# Functional recruitment of astrocyte-derived neurons into the mouse visual cortex

**DOI:** 10.64898/2026.01.25.701593

**Authors:** Sydney Leaman, Filipe Machado Ferreira, Lin Dong, Ana Beltrán Arranz, Jerónimo Jurado-Arjona, Araceli García Mora, Alexis Cooper, Nina Treder, Syauqina Fakhira, Varun Sreenivasan, Nicolás Marichal, Oscar Marín, Adil Khan, Benedikt Berninger

**Affiliations:** Centre for Developmental Neurobiology, Institute of Psychiatry, Psychology & Neuroscience, King’s College London, UK; Medical Research Council Centre for Neurodevelopmental Disorders, Institute of Psychiatry, Psychology & Neuroscience, King’s College London, London, UK; Institute of Physiological Chemistry, University Medical Center Johannes Gutenberg University, Mainz, Germany; The Francis Crick Institute, London, UK

**Author notes:** These authors contributed equally (Adil Khan, Benedikt Berninger).

## Abstract

Inducing neurogenesis from glia could permit circuit remodeling and neuron replacement in the brain. Indirect evidence suggests that glia-to-neuron reprogramming is feasible, but the dynamics of conversion have not been directly observed, and it remains unclear whether glia-derived cortical neurons can functionally integrate into brain circuits. Here, we use two-photon imaging to visualize the reprogramming of astroglia into induced neurons in the mouse cortex *in vivo*. We track the emergence of spontaneous neuron-like calcium transients in astrocyte-derived induced neurons, demonstrating their functional activity *in vivo*. Importantly, these induced neurons exhibit orientation tuned, visually evoked calcium responses, demonstrating their functional integration into cortical circuitry - a prerequisite for their utility in circuit repair. Thus, astrocytes can be recruited to generate functional neurons in the postnatal cortex, enabling circuit remodeling in perinatal injury or neurodevelopmental disorders without relying on external cellular sources.

## Introduction

In placental mammals, neurogenesis is largely completed *in utero*(*1, 2*), leaving us neurologically vulnerable to perinatal disease or injury affecting cortical neurons. Strategies to reactivate neurogenesis in the neocortex might have therapeutic potential, provided that newly generated neurons demonstrate functional activity and are appropriately recruited into existing cortical circuits. Although the postnatal mammalian neocortex lacks a dedicated neural stem cell population, one approach would be to engineer cortical neurogenesis *in situ* by reprogramming local non-neuronal cell populations. Astrocytes are ubiquitous in the cortex, share a close developmental origin with neurons(*3–5*) and, in early postnatal life, are still generated via local proliferation(*6, 7*). Thus, astrocytes are a promising potential cellular source for *in vivo* lineage reprogramming and circuit remodeling.

Pioneering studies demonstrated that early postnatal cortical astrocytes *in vitro* could be reprogrammed into neurons, capable of generating action potentials and forming functional synapses, following forced expression of transcription factors involved in neuron specification such as Ascl1, Dlx2 and Neurog2(*8–10*). Injury can elicit the proliferation of cortical macroglia (reactive astrocytes and oligodendrocyte progenitor cells)(*11–14*). Several studies have targeted these cells with retroviral and, more recently, transgenic approaches to express neurogenic transcription factors, inducing neuronal conversion *in vivo*(*14–18*). Recently, we demonstrated lineage conversion of astrocytes into induced neurons (iNs), exhibiting hallmarks of cortical interneurons, in the uninjured postnatal mouse cortex (*19, 20*). Glia have also been used as a cellular source to experimentally induce neurogenesis in several other regions of the central nervous system(*21–26*). However, the evidence for lineage conversion in these studies has been indirect.

Excitingly, several studies report that iNs show electrophysiological evidence of synaptic inputs in *ex vivo* tissue preparations(*17, 19, 21, 22, 24*). Additionally, circuit tracing techniques in the hippocampus and striatum have demonstrated structural synaptic inputs onto iNs(*21, 22*). Together, these findings suggest that iNs generated *in vivo* receive some synaptic input, but whether they integrate into functional circuits and participate during sensory processing in awake animals remains unknown.

Claims have been made that adult mouse cortical astrocytes could be reprogrammed into iNs, showing remarkable structural and functional fidelity to fully mature cortical neurons *in vivo.* However, subsequent results have confirmed that the AAV vectors used in these studies were, in fact, labelling pre-existing endogenous neurons rather than mediating lineage conversion(*27–29*). These missteps emphasize the importance of directly tracking *in vivo* lineage reprogramming(*30*) and establishing the nature and timing of cellular events during conversion.

In this work, we tracked the dynamic cellular morphology and the emergence of spontaneous calcium activity in nascent astrocyte-derived neurons induced by retroviral expression of Ascl1SA6(*31, 32*) and Dlx2 – two critical regulators of cortical interneuron fates(*33, 34*). After demonstrating their structural maturation and continued functional activity after eye-opening, we further demonstrate that Ascl1SA6-Dlx2 iNs in the primary visual cortex (V1) exhibit visually evoked calcium responses, tuned to orientation of grating stimuli in awake animals. Together, these results show that early postnatal astrocytes can serve as a cellular source for generating functional neurons that integrate into mammalian cortical circuitry.

### Direct visualization of induced neurogenesis in the postnatal mammalian cortex

Whilst naturally occurring neurogenesis concludes prenatally in the mouse cortex, gliogenesis is ongoing, with progenitors of astrocyte and oligodendrocyte lineages proliferating locally during the first postnatal week(s)(*6, 7*). To selectively express proneural transcription factors Ascl1SA6(*19, 31, 32*) and Dlx2 (and a red fluorescent protein, RFP) in cortical glia we injected a γ-retroviral vector, which relies on mitosis for nuclear entry and subsequent genome integration(*35, 36*), into the mouse cortex at postnatal day 5-6 (P5-6). We implanted a cranial window overlying the viral injection site and used two-photon imaging to visualize the morphology of transduced cells over the following days (**Fig. 1A**). Viral RFP expression was detectable from around 38-42 hours post-injection ( hpi), gradually increasing over a day or so (**Fig. S1A**). Initially, transduced cells showed glial-like morphologies, with many short, hair-like processes emanating from a central soma and many examples of transduced cells ensheathing blood vessels, a characteristic of astroglia (**Fig. 1B-C, Fig. S1C**, green arrows). Over the subsequent days, transduced cells underwent rapid and dramatic morphological changes (**Fig. 1C-E**, **Fig. S1**). We observed individual glia at 48 hpi with short, bushy glial-like processes, proliferate and give rise to daughter cells that each extended long, neurite-like processes and appeared to migrate short distances by 4 days post-injection ( dpi; **Fig. 1D**). Morphological reconstructions of Ascl1SA6-Dlx2 transduced cells (**Fig. 1E-F**) from several (n=7) mice imaged between 2-5 dpi, revealed a simplification of cellular branching structure, indicated by a reduction in both the number of primary processes emanating from the soma and total count of terminal endpoints of all processes within each cell (**Fig. 1G-H**). Meanwhile, remaining processes showed marked increase in length over time, with the average branch length increasing by approximately 18 μm per day, and corresponding increases in the longest-shortest path and size of the convex hull occupied by the cells (**Fig. 1I-K, Table S1**). By 4 dpi, most cells showed simple unipolar or bipolar morphologies (65.8%, 90/146, no. primary process ≤2) with the longest processes measuring, on average, over 100 μm (mean 114 μm, SD 66.3 μm), consistent with conversion of glia towards immature neurons. In contrast, cortex injected with a control virus encoding RFP-only (MMLV-CAG-IRES-RFP) revealed RFP+ cells with glial morphologies at 4 dpi, most of which expressed astrocyte marker Sox9 (**Fig. S2,** mean 79.04%, 95%CI [70.86%, 85.39%]), in keeping with previous reports(*6, 37*).

**Fig. 1.**
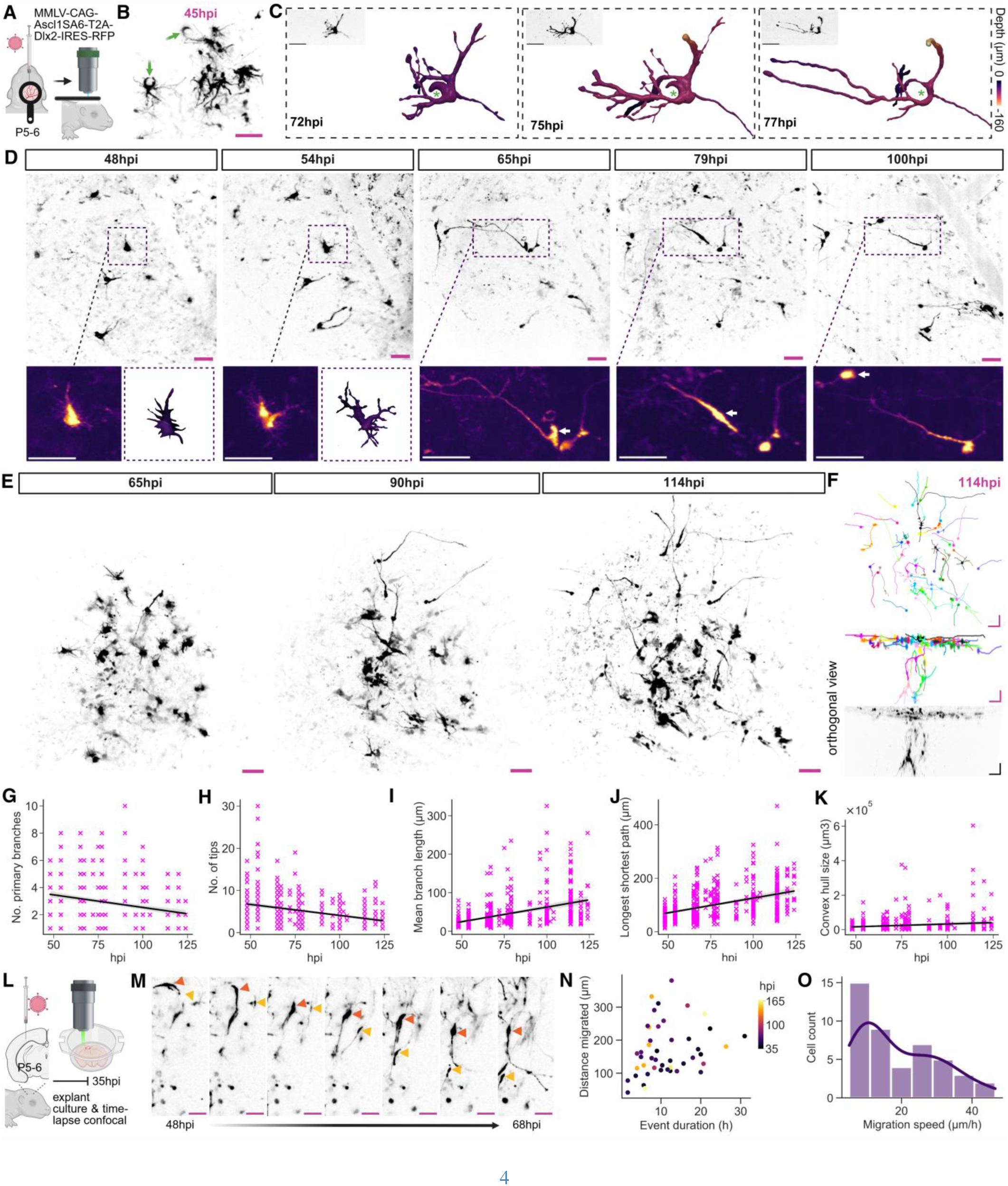
Direct visualization of Ascl1SA6-Dlx2 induced neurogenesis from cortical glia. (**A**) Experimental schematic. **(B)** Example of transduced glia wrapped around blood vessels at 45 hpi. **(C)** Rendered examples of a transduced cell expressing Ascl1SA6-Dlx2-RFP ensheathing a blood vessel (green asterisk) which extends processes and then simplifies its branching structure, loosening contact with the blood vessel by 77 hpi. MIP of raw image in top left inset. Scale bars: 25 μm. **(D)** MIP from two-photon imaging showing an example of a glial cell proliferating and giving rise to two daughter cells (48-65 hpi) that develop long neurite-like processes, with one migrating a short distance (white arrows in lower panels). **(E)** MIP of representative imaging site from 65-114 hpi. Full time series of the same imaging site from 54-120 hpi can be found in **Fig. S1**. **(F)** Example of individual cell reconstructions of imaging site shown in (E) at 114 hpi (top panel: top view, middle and bottom panel: orthogonal view of reconstructions and orthogonal MIP, respectively). Colored lines indicate individual cells with circles indicating soma locations. **(G-K)** Relationship between named morphological measurements and time in hours post-injection ( hpi). See Table S1 for statistical analysis. **(L)** Schematic of experiment for data shown in (M-O). **(M)** Cropped MIP example from time-lapse confocal microscopy of an explant culture of a coronal cortical slice injected with Ascl1SA6-Dlx2-RFP from 48-68 hpi. Yellow and purple arrows indicate examples of two RFP+ cells with neuronal-like morphology that migrate during this time. **(N-O)** Quantifications of the distance, duration and speed of migration events in Ascl1SA6-Dlx2 transduced cells with neuronal-like morphologies (see material and methods). Scale bars: 50um, unless otherwise indicated.

Migration is an important behavior of all newborn neurons, allowing them to locate themselves appropriately within tissue layers and structures. Cortical interneurons need to migrate long distances from their birthplace in the ganglionic eminences into the developing dorsal telencephalon, a process partly orchestrated by Ascl1 and Dlx2(*33, 38–40*). We observed individual Ascl1SA6-Dlx2 transduced cells with morphological features of immature neurons that changed positions relative to other cells and anatomical landmarks within our field of view, as well as a general spatial spread of labelled cells over time (**Fig. 1D-E**). While consistent with migration, the temporal resolution of our acquired data was limited in live pre-weaned pups.

Thus, to complement our *in vivo* imaging, we performed time-lapse confocal microscopy on explant cultures of coronal brain slices of cortex injected with MMLV-CAG-Ascl1SA6-T2A-Dlx2-RFP (Ascl1SA6-Dlx2-RFP) at P5-6 (**Fig. 1L-O**, see **Methods**). This allowed image acquisition at 20-minute intervals over several days from the onset of viral expression around 35-40 hpi (**Movie S1**). We observed many individual cells undergoing migration events from one stationary location to another during the imaging period. Cells that migrated covered distances up to several hundred microns (**Fig. 1M-N**; range 41.44 - 380.87 μm, 45 cells from n=4 mice) at mean speeds of approximately 20 μm/hour (mean = 19.23 μm/h, SD 11.34; **Fig. 1O**) and displayed immature neuron-like morphologies (**Movie S1**), confirming migratory behavior of putative nascent induced neurons (iNs). In summary, cortical glia expressing Ascl1SA6 and Dlx2 exhibit morphological changes and migratory behavior consistent with conversion to immature neuronal identities within 4-5 days from viral injection.

Electrical and calcium activity are necessary components during neuronal development, regulating differentiation, migration, and survival of newborn neurons(*33, 41–43*). By 4-5 dpi most Ascl1SA6-Dlx2 transduced glia had acquired immature neuronal morphologies, so we next asked if these structural changes coincided with a divergence from astrocyte-like calcium dynamics, which are dominated by slow, highly correlated calcium waves(*44, 45*). To accomplish this, we performed injections of Ascl1SA6-Dlx2-RFP into the cortex of transgenic mice that expressed cre-dependent GCaMP6s in cortical astrocytes (mGFap-Cre;Ai96(RCL:GCaMP6s), **Fig. 2**), allowing us to simultaneously record calcium activity in local astrocytes and Ascl1SA6-Dlx2 iNs originating from astrocytes. As expected, local cortical astrocytes that were not transduced by the retrovirus (i.e. RFP-, GCaMP6s+ astrocytes) had infrequent, synchronized calcium events (**Fig. 2B, D -** green, **Table 2**). At 48 hpi, most Ascl1SA6-Dlx2 transduced cells showed near-absent or more infrequent calcium events (**Fig. 2B, D -** magenta), albeit not statistically significantly different to their astrocytic counterparts at this timepoint (p = 0.31, see **Table 2**). From around 4 dpi (96 hpi) onwards, however, Ascl1SA6-Dlx2 transduced cells showed clear spontaneous calcium transients (**Fig. 2C-D, Movie S2**), with calcium event frequency in Ascl1SA6-Dlx2 transduced cells increasing over time (estimated increase of ∼0.5events min^-1^ per day post-injection, *p* < 0.001, see **Methods**, **Table 2**). Meanwhile, non-transduced astrocytes did not meaningfully differ in the frequency of calcium events over the same period (+0.0003 events min⁻¹ hour^-1^, *p* = 0.53, **Table 2**, **Fig. 2D, Table S2).** Examining pairwise correlations between non-transduced astrocytes and Ascl1SA6-Dlx2 iNs within the same recording session at timepoints after 96 hpi (4 dpi) revealed decorrelation of iNs from the relatively synchronous calcium activity patterns observed in non-transduced astrocyte pairs (**Fig. 2E**). These findings, in conjunction with the visualized morphological changes (**Fig. 1, Fig. S1, Fig 2.B-C**), are in keeping with the rapid conversion of glia into immature neurons within a week of Ascl1SA6-Dlx2 expression.

**Fig. 2.**
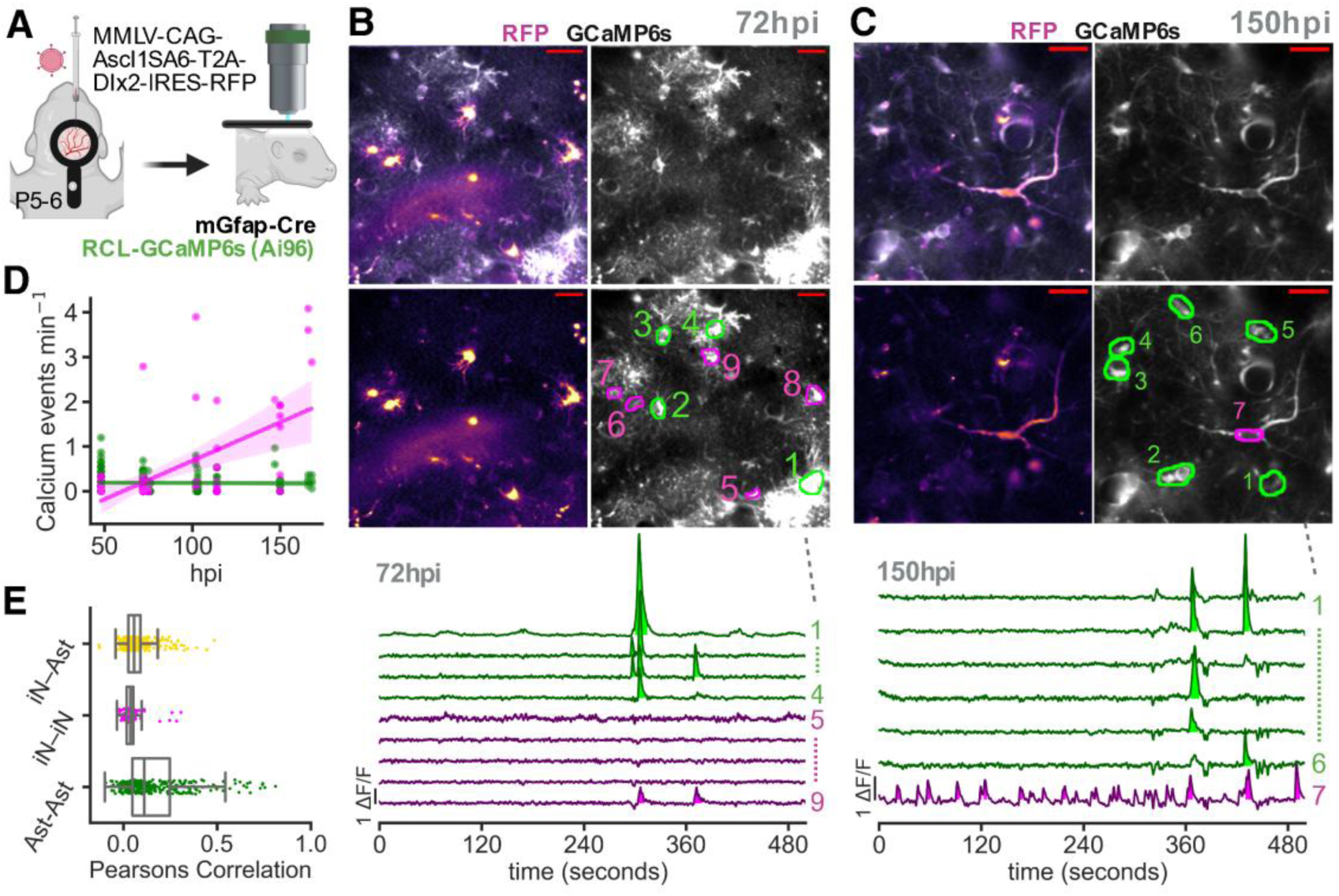
Emergence of early neuron-like calcium transients in Ascl1SA6-Dlx2 transduced cells. **(A)** Schematic of experimental design. **(B)** Top: mean intensity projection of example imaging site at 72 hpi showing transduced cells (magenta) and surrounding non-transduced cells (green numbered regions of interest; ROIs) expressing GCaMP6s. Bottom: calcium activity snippet recorded from cells shown in top of (B). Green traces illustrate data for non-transduced local astrocytes, magenta traces indicate Ascl1SA6-Dlx2 transduced cells (RFP+). **(C)** Top: mean intensity projection of imaging site at 150 hpi showing example of Ascl1SA6-Dlx2 transduced cell (magenta ROIs) and surrounding non-transduced astrocytes (green ROIs) expressing GCaMP6s. Bottom: calcium activity snippet recorded from cells shown in Top of (C). **(D)** Relationship between time ( hpi) and frequency of calcium events in Ascl1SA6-Dlx2 transduced (RFP+) cells (magenta) and nearby GCaMP6s+ astrocytes (green). See Table S2. **(E)** Pearsons correlation (y-axis) between pairs of cells recorded from later than 96 hpi (4 dpi) from the same recordings. Pairwise relationships are categorized as between two Ascl1SA6-Dlx2 transduced putative induced neurons (iN-iN), between two non-transduced GCaMP6s astrocytes (Ast-Ast), or between one iN and one GCaMP6s+ astrocyte (iN-Ast). One-way ANOVA, p<0.0001. Pairwise Tukey: Ast-Ast vs. iN-Ast mean difference = −0.093, 95%CI [-0.118, - 0.068], p<0.0001; Ast-Ast vs. iN-iN mean difference = −0.123, 95% CI [-0.158, −0.088], p<0.0001; iN-Ast vs. iN-iN mean difference = −0.030, 95%CI [-0.065, 0.005], p=0.11 (non-significant). Scale bars: 50um.

### Structural maturation of astrocyte-derived Ascl1SA6-Dlx2 interneuron-like iNs

To ultimately assess whether Ascl1SA6-Dlx2 iNs could be recruited into local circuitry, we selected V1 as an experimentally accessible region with well characterized functional response and feature selectivity properties. We waited until after eye-opening and then performed cranial windows on mice who had previously received an injection of MMLV-Ascl1SA6-T2A-Dlx2-IRES-RFP into visual cortex at P5-6 (**Fig. 3A**). Whilst structural imaging volumes acquired at 13-14 dpi revealed that RFP-only control virus transduced cells showed glial morphology (**Fig. 3B**), Ascl1SA6-Dlx2 expressing cells exhibited neuronal morphologies, appearing more mature than those we characterized during the first week of reprogramming (**Fig. 3B-C, Movie S3**). Comparing Ascl1SA6-Dlx2 iNs at 13-14 dpi to those we had previously characterized at 4-6 dpi, we found they had more primary processes and longer mean branch length (n=6 mice, 92 cell reconstructions at 13-14 dpi; **Fig. 3D-E**).

**Fig. 3.**
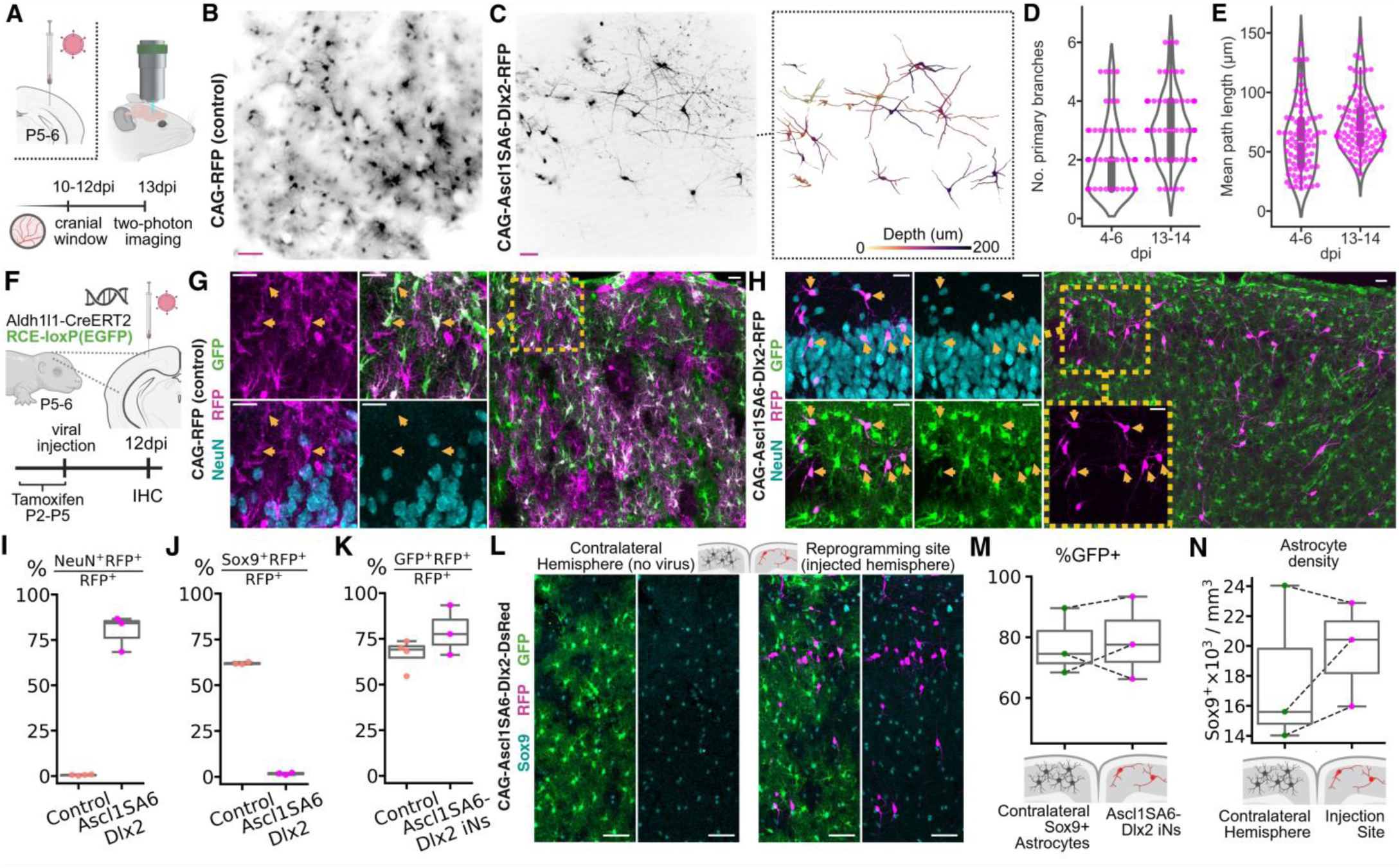
Structural maturation and lineage origins of Ascl1SA6-Dlx2 iNs. **(A)** Experimental schematic of two-photon imaging of Ascl1SA6-Dlx2 iNs at 2 wpi. **(B)** Representative MIP of two-photon structural stack acquired at 1060nm wavelength of MMLV-CAG-IRES-RFP control cells showing astrocyte-like morphologies. **(C)** Representative MIP of two-photon structural stack acquired at 1060nm wavelength of MMLV-CAG-Ascl1SA6-T2A-Dlx2-IRES-RFP control cells showing neuronal morphologies (left). Right: projection of 3D rendered cell structures color scale indicating depth from most superficially located rendered cell (upper layer 1). **(D)** Number of primary branches at 4-6 dpi compared to 13-14 dpi of Ascl1SA5-Dlx2 iNs. The unpaired mean difference between groups is 1.14 primary branches, 95.0%CI [0.793, 1.47]. **(E)** Comparison of mean branch length in Ascl1SA6-Dlx2 iN reconstructions where unpaired mean difference between 4-6 dpi and 13-14 dpi is 20.5 μm 95.0%CI [7.12, 36.6]. **(F)** Schematic of experimental design for genetic labelling of astrocyte lineage prior to retroviral injection. **(G)** MIP of coronal sections of cortex injected with MMLV-CAG-IRES-RFP control retrovirus at 12 dpi. Yellow arrows indicated examples of RFP+ transduced cells (magenta) that show glial morphologies, lack NeuN expression (cyan) and express GFP (Aldh1l1-CreERRT2;RCEloxP(GFP); green). Scale bar = 25 μm. **(H)** MIP of coronal section of cortex injected with MMLV-CAG-Ascl1SA6-T2A-Dlx2-IRES-RFP retrovirus at 12 dpi. Yellow arrows indicated examples of RFP+ transduced cells (magenta) that show neuronal morphologies, express both NeuN (cyan) and GFP (Aldh1l1-CreERRT2;RCEloxP(GFP); green) **(I-K)** Percentage of the population of transduced (RFP+) cells that co-express NeuN (D), Sox9 (E) and GFP (F) in RFP-only control vs. Ascl1SA6-Dlx2 injected cortices. **(L)** Representative example of Sox9 (cyan) and GFP (green) expression in the non-injected contralateral hemisphere (left) and the same in the hemisphere injected with Ascl1SA6-Dlx2 (magenta). No no loss of astrocyte density as indicated by GFP distribution and density of Sox9 (quantified in (N)). **(M)** Quantification of the percentage of all cells in the groupings indicated on the x-axis that co-express GFP. Left (green dots) quantifies the percentage of Sox9+ cells in the contralateral (i.e. non-injected) hemisphere that express GFP. Right (magenta dots) show the percentage of Ascl1SA6-Dlx2 iNs (RFP+NeuN+ cells) that express GFP (see images in H, L). Dashed lines connect data points collected from an individual animal. Paired mean difference of 1.55%, 095%CI [-8.33, 7.39], p=0.53 two-side permutation test. **(N)** Quantification of the density of astrocytes (Sox9+ cells) at the Ascl1SA6-Dlx2 injection site (right, magenta dots) and density at a comparable anatomical location in the contralateral (non-injected) hemisphere (left, green dots). Dashed lines connect data points collected from an individual animal. Paired mean difference 1.86×10^3^ mm^-3^, 95.0%CI [-1.16, 3.85 x10^3^mm^-3^], p=0.505 two-side permutation test.

Since it was not possible to maintain headplates and cranial windows following viral injection at P5-6 for more than a week, we used an independent experimental approach to confirm the lineage identity of Ascl1SA6-Dlx2 iNs that we observed in live imaging at 2 wpi. We genetically labelled astrocytes with GFP, permanently and with high specificity, by administering tamoxifen to Aldh1l1-CreERT2;RCE:loxP (Cre-dependent GFP) mice from P2-P5 (**Fig. S3**). We then injected either RFP-only (control) or Ascl1SA-Dlx2-RFP retroviral vectors into V1 at P5-6 and collected cortical tissue at 12 dpi (**Fig. 3F**).

As expected, in the RFP control-injected cortex, transduced cells showed glial morphologies with most expressing astrocyte marker Sox9 (61.85%, 95%CI [61.15%, 62.55%]) and lacking expression of the neuronal marker, NeuN (Rbfox3, **Fig. 3G, I-J**). In sharp contrast, nearly all RFP+ cells in Ascl1SA6-Dlx2 injected cortices showed neuronal morphologies (**Fig. 3H**) with most expressing NeuN (80.76%, 95%CI [68.43%, 89.05%], **Fig. 3I)** and only very few remaining Sox9 positive (1.54%, 95%CI [0.96%, 2.47%], **Fig. 3J**).

Of all Ascl1SA6-Dlx2 induced neurons (i.e. RFP+, NeuN+ cells), the majority (82.03%, 95%CI [59.13%, 93.51%]) expressed GFP (**Fig. 3H, K**), indicating they arose from Aldh1l-CreERT2 expressing astrocyte lineage origins. A slightly lower fraction of RFP-only control virus transduced cells expressed GFP (66.92%, 95%CI [58.53%, 74.36%], **Fig. 3K**), which may be explained by proliferation or survival of transduced oligodendrocyte precursor cells (OPCs) in the absence of reprogramming factors (although the difference between groups was not statistically significant, unpaired mean difference 12.4% (95%CI [-1.3, 27.3], p=0.196 two-side permutation test, **Fig. 3K**). The percentage of GFP-expressing Ascl1SA6-Dlx2 induced neurons was very similar to the percentage of Sox9+ astrocytes in the contralateral (non-injected) hemisphere that were genetically labeled by GFP in our experiment (paired mean difference of 1.55%, 95%CI [-8.33, 7.39], p=0.53 two-side permutation test, **Fig. 3L-M**). These data confirmed that the Aldh1l1+ astrocyte lineage is the predominant source of Ascl1SA6-Dlx2 iNs.

Notably, we did not observe a reduction of overall astrocyte density at the Ascl1SA6-Dlx2 cortical injection site at 12 dpi as compared to the contralateral (non-injected) hemisphere (**Fig. 3L,N**). This suggests that in this developmental context (where astrogenesis is ongoing at the time of viral injection), astrocytes can maintain their numbers despite ensuing neuronal lineage reprogramming.

Given our previous findings that Ascl1SA6 (in combination with Bcl2) generated neurons with hallmarks of GABAergic neuronal identities and the established role of Dlx2 and Ascl1 in acquisition of cortical inhibitory neuron identity, we hypothesized that astrocyte-derived Ascl1SA6-Dlx2 iNs might acquire features of inhibitory neuron identity. We injected MMLV-CAG-Ascl1SA6-T2A-Dlx2-IRES-RFP into the P5-6 cortex of Vgat-ires-Cre:RCE:loxP(GFP) and Nex(Neurod6)-Cre;RCE:loxP(GFP) transgenic mice, which drive cre-recombinase expression (and subsequent GFP reporter expression) in pan-inhibitory and -excitatory neuronal populations, respectively (**Fig. S4A**). In Nex(Neurod6)-Cre;RCE:loxP(GFP) mouse cortex, which labels most neurons in the mouse cortex, Ascl1SA6-Dlx2 iNs did not co-localize with GFP expression (**Fig. S4B-C**). Thus, Ascl1SA6-Dlx2 iNs are not generating glutamatergic-like phenotypes at this timepoint and cannot be explained by spuriously labelled pre-existing cortical excitatory neuron populations. In contrast, in Vgat-Cre:GFP cortices, around a third of Ascl1SA6-Dlx2 transduced cells expressed GFP (36.93%, 95%CI [28.55%, 46.17%]) and a similar proportion co-expressed GABA (37.27%, 95%CI [26.76%, 49.12%]) (**Fig. S4D-E**), indicating that Ascl1SA6-Dlx2 iNs can acquire interneuron-like molecular phenotypes (**Fig. S4**).

### Spontaneous activity and functional recruitment of Ascl1SA6-Dlx2 iNs into cortical circuits

To ascertain whether Ascl1SA6-Dlx2 iNs are functionally active *in vivo* at 13-14 dpi, we recorded spontaneous calcium activity from RFP+ iNs, along with neighboring (non-transduced) GCaMP6s+ astrocytes, in awake mice on a stable platform in the dark at 13 dpi (**Fig. 4A**). Immunohistochemistry in *ex vivo* tissue collected at 16 dpi revealed that the majority of Ascl1SA6-Dlx2 transduced (RFP+) cells expressed NeuN (77.56%, 95%CI [63.95%, 87.07%]), with around one fifth of these iNs (21.44%, 95%CI [8.93%, 43.18%]) expressing GCaMP6s (**Fig. S5**). Ascl1SA6-Dlx2 iNs showed spontaneous calcium activity that was distinct from the patterns of activity in surrounding GCaMP6s+ astrocytes, with calcium events occurring at greater frequency and having shorter mean duration and lower amplitudes than their astrocyte counterparts (**Fig. 4B-C, E-G, Movie S4**). Pairwise correlations between individual cells in the same recording revealed that Ascl1SA6-Dlx2 iNs activity is decorrelated from the highly correlated activity between astrocyte pairs (**Fig. 4C-D, H**). This would be in keeping with distinct drivers of functional activity in Ascl1SA6-Dlx2 iNs as compared to their astrocyte counterparts. With a retrovirus expressing only RFP (MMLV-CAG-IRES-RFP), transduced cells expressing GCaMP6s had distinctly astrocytic morphology and patterns of calcium activity that closely resembled that of nearby non-transduced astrocytes (**Fig. S6**). Thus, at 2 wpi, Ascl1SA6-Dlx2 iNs exhibit spontaneous neuron-like functional activity *in vivo* in the awake mouse cortex.

**Fig. 4.**
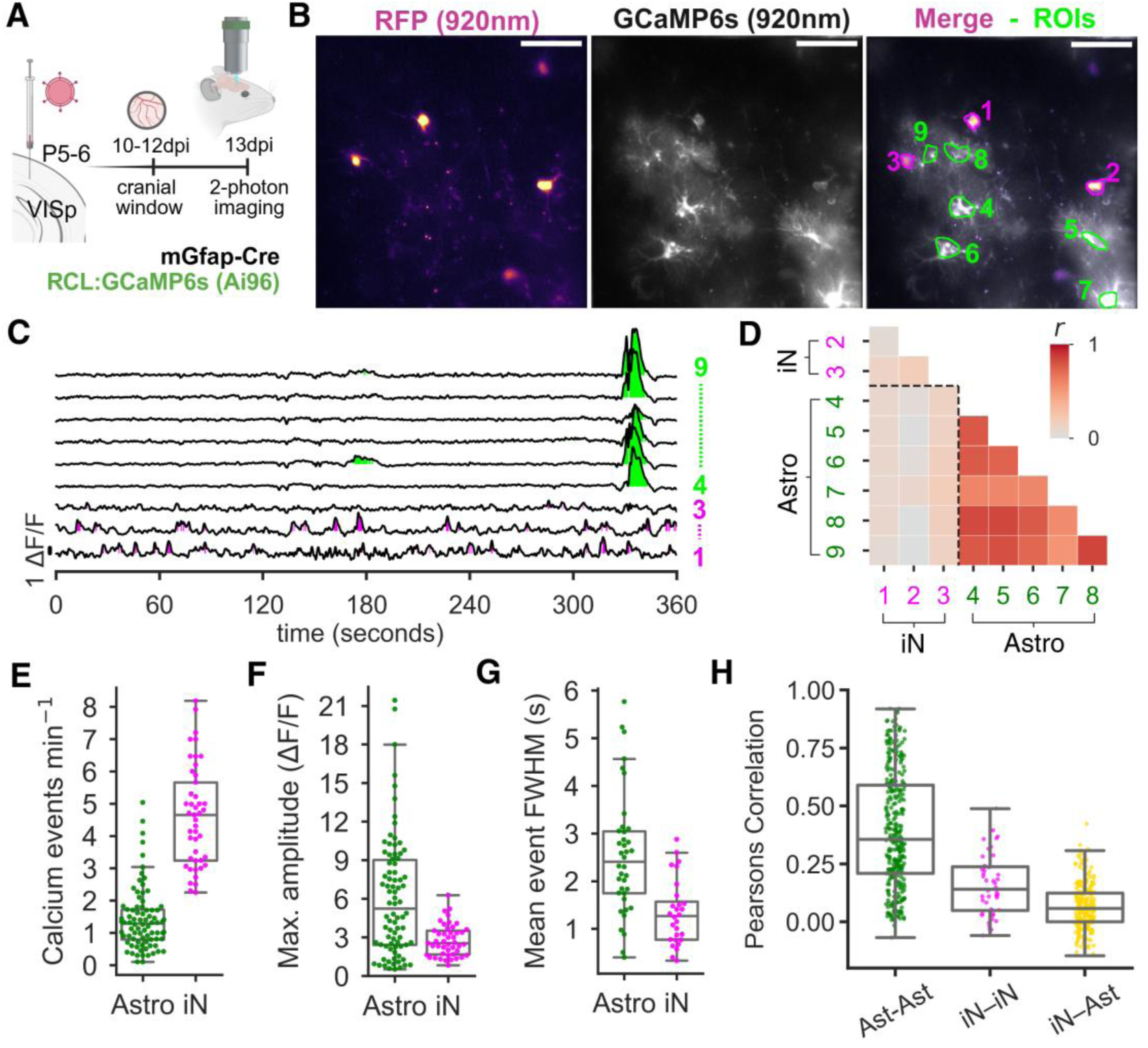
Ascl1SA6-Dlx2 iNs show spontaneous *in vivo* calcium activity at 2 wpi. **(A)** Schematic of experimental design. **(B).** Mean intensity projection of red (RFP) and green (GCaMP6s) channels from representative functional recording session acquired at 920nm excitation wavelength with Ascl1SA6-Dlx2 iN (RFP+ cells) regions of interest (ROI) shown in magenta and non-transduced GCaMP6s astrocyte ROI in green. Scale bar = 50 μm. **(C)** Example of spontaneous calcium activity (ΔF/F) recorded from cells labelled in (B). **(D)** Matrix showing Pearson’s correlation (r) between pairs of cells labelled in (B) and (C). **(E)** Comparison of calcium event frequency between astrocytes and Ascl1SA6-Dlx2 iNs in entire dataset (unpaired mean difference: 3.2 events per min, 95%CI [2.71, 3.7]). **(F)** Comparison of calcium event maximum amplitudes between astrocytes and Ascl1SA6-Dlx2 iNs (unpaired mean difference: −3.33 ΔF/F, 95%CI [-4.49, −2.32]). **(G)** Comparison of calcium event durations (mean FWHM, seconds) between astrocytes and Ascl1SA6-Dlx2 iNs (unpaired mean difference: −1.21 seconds, 95%CI [-1.7, −0.753]). N=5 mice (45 iNs, 82 astrocytes) for data in (E)-(G). **(H)** Pearsons correlation (y-axis) between: pairs of Ascl1SA6-Dlx2 iNs (iN-iN), two non-transduced GCaMP6s astrocytes (Ast-Ast), or one iN and one GCaMP6s+ astrocyte (iN-Ast). One-way ANOVA, p= <0.0001. Pairwise Tukey: Ast-Ast vs. iN-Ast mean difference = −0.328, 95%CI [-0.367, - 0.289], p<0.0001; Ast-Ast vs. iN-iN mean difference = −0.234, 95%CI [-0.306, −0.162], p<0.0001; iN-Ast vs. iN-iN mean difference = 0.094, 95%CI [0.020, 0.168], p=0.008.

Our previous work showed that iNs derived from astrocytes in the early postnatal cortex of mice show spontaneous excitatory post-synaptic currents in *ex vivo* patch clamp electrophysiology(*19*). This suggests they receive excitatory pre-synaptic inputs. However, it is unknown whether iNs can be functionally recruited during sensory perception in awake animals. Having established a model to observe calcium transients as a readout of their functional activity in V1 at 2 wpi, we next asked whether Ascl1SA6-Dlx2 iNs show evoked responses to visual stimuli presented to the mice.

We presented the mice drifting visual gratings of two second duration and eight, randomly interleaved orientations (or blank, grey screen) whilst recording GCaMP6s activity from RFP+ cells in Ascl1SA6-Dlx2 or RFP-only control injected cortices (**Fig. 5A**). Transduced cells in RFP-only control virus injected cortices that co-expressed GCaMP6s exhibited astrocyte-like calcium transients (**Fig. 5B-C**). Stimulus-onset aligned, trial-averaged responses of RFP-only control astrocytes did not reveal evoked calcium responses following presentation of visual stimuli (**Fig. 5D, H-K,** 2.86%, 2/70 cells, pooled from n=4 mice showed significant responses to at least one stimulus, see **Methods**). Similarly, non-transduced GCaMP6s+ astrocytes in the same field of view as Ascl1SA6-Dlx2 iNs did not show visual responses (**Fig. 5E-K**; 0.50%, 1/202, responsive cells pooled from n=9 mice).

**Fig. 5.**
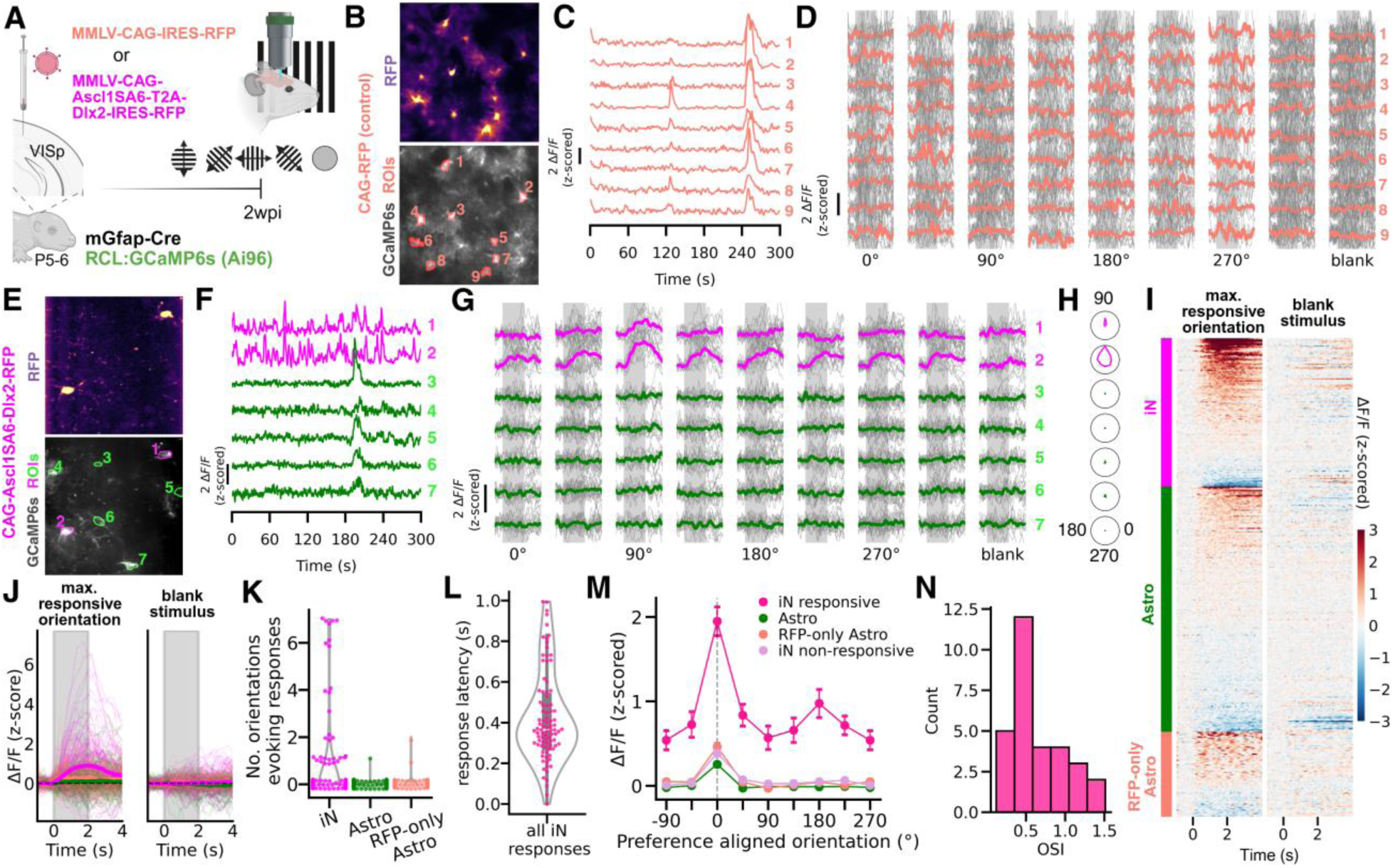
Functional integration of Ascl1SA6-Dlx2 iNs into V1. **(A)** Experimental schematic. **(B)** Mean image acquired from a functional recording at 920nm wavelength following RFP-only control virus injected into mGfap-Cre;Ai96(GCaMP6s) mouse cortex. Numbered ROIs identify corresponding calcium activity and responses similarly numbered in (C)-(D). **(C)** Example of calcium activity recorded from numbered ROIs identified in (B) during visual stimulus presentation. RFP-only control virus transduced GCaMP6s+ astrocytes show typical astrocyte-like responses with highly synchronized, slow, infrequent calcium events. **(D)** Stimulus aligned responses separated into the eight different orientations of drifting gratins presented and gray screen (blank) stimulus. Vertical bars shaded in gray indicate the timing of the stimulus presentation (2s). Thin gray traces show ΔF/F calcium responses for individual trials. Thick colored (salmon) trace shows the mean of all trials to each visual orientation/blank. (**E-G**) As for (B)-(G) but for Ascl1SA6-Dlx2-RFP injected cortex. Magenta ROIs and traces indicate Ascl1SA6-Dlx2 transduced cells. Green ROIs and traces indicated GCaMP6s+ astrocytes from the same field of view as the injection site that were not transduced by the virus. (E) shows MIP of mean images from two simultaneously recorded planes 10 μm apart. **(H)** Polar plots as a visual summary of cells tuning properties for each cell numbered in (E)-(G) normalized to the highest magnitude response for this recording. **(I)** Raster plot showing the trial-averaged, stimulus aligned response for: the grating orientation resulting in the greatest magnitude response for each cell (left); and the blank (gray-screen) stimulus (right). Cells sorted first by grouping (Ascl1SA6-Dlx2 iN (magenta, iN, top), GCaMP6s+ non-transduced astrocyte in (green, Astro, middle), RFP-control transduced astrocyte (salmon, RFP-only Astro, bottom)) then by maximum magnitude responses (between stimulus onset and 500ms following stimulus offset). **(J)** As for (I) but with z-scored ΔF/F trial-averaged, stimulus aligned responses represented on the y-axis with individual cell traces presented as thin lines using same color scheme as previous panels to denote cell groupings. Thicker bold lines represent the mean response of all cells within each grouping. **(K)** Graph showing the total number of grating orientations for which a given cell (dots) showed a statistically significant evoked calcium response (see materials and methods). Ascl1SA6-Dlx2 iNs (iN, magenta), GCaMP6s+RFP- astrocytes (Astro, green), RFP-only control virus transduced astrocytes (RFP-only Astro, salmon). **(L)** Response latencies (defined as time taken to reach 20% of maximum response magnitude) for all significant responses of Ascl1SA6-Dlx2 iNs to any orientation. **(M)** Response magnitudes (z-scored ΔF/F) for each orientation of visual gratings with preferred orientations for each cell aligned to 0 degrees. Dots indicate mean magnitude, bars show ± SEM. Ascl1SA6-Dlx2 iNs were divided into two groups based on whether a cell showed statistically significant visual responses to any orientation (“iN responsive”, deep pink hue), or not (“iN non-responsive”, pale purple). **(N)** Orientation selectivity index (OSI) of Ascl1SA6-Dlx2 iNs that showed visual responses to at least one orientation. Values >1 result from negative signed calcium responses at orientations orthogonal to the preferred stimulus.

In contrast, around a quarter of Ascl1SA6-Dlx2 iNs recorded from (24.59%, 30/122 cells pooled from N=9 mice) showed significant visually evoked calcium responses during presentation of at least one orientation of the visual grating stimulus (**Fig. 5E-K, Movie S5**). More than half (53.33%) of these iNs showed responses to multiple orientations (**Fig. 5K**) with latency of visual responses of around 400ms (median 363ms, mean 423ms, SD 220ms, defined as time taken to reach 20% of maximal response, **Fig. 5L**). Visually evoked calcium responses peaked, on average, towards the end of the 2 second stimulus presentation (mean 2.04s, SD 0.52s). We aligned responses to each cells preferred orientation (i.e. the orientation of the grating stimulus which elicited the greatest magnitude response for each cell) and observed that Ascl1SA6-Dlx2 iNs showed a second peak of response magnitudes at 180 degrees to the preferred stimulus (p=0.005, one-sided, one-sample t-test, **Table S3,** see **Methods**), indicating some tuning to orientation (**Fig. M-N**). Thus, retroviral-mediated expression of Ascl1SA6 and Dlx2 in proliferating mouse cortical astrocyte progenitors can generate neurons capable of functional recruitment into sensory circuits *in vivo*.

## Discussion

In this study, we used *in vivo* two-photon imaging to directly visualize the conversion of early postnatal cortical glia into neurons. We showed that iNs were derived from fate-committed postnatal astrocytes and exhibited features of inhibitory, but not glutamatergic, identity. We observed the emergence of spontaneous neuron-like activity in iNs and their functional decoupling from the highly synchronized activity of astrocyte networks, suggesting iNs develop distinct drivers of activity from their astrocytic origins. Finally, we discovered that iNs show visually evoked calcium responses tuned to orientation, confirming their recruitment into sensory cortical circuitry.

Our longitudinal imaging in upper cortical layers revealed a highly dynamic process of morphological remodeling in Ascl1SA6-Dlx2 transduced cells taking place *in vivo*. These data not only provide important evidence to demonstrate the authenticity of induced neurogenesis in the postnatal cortex but also reveal a surprisingly rapid timescale for conversion. Most Ascl1SA6-Dlx2 transduced cells showed features of immature neurons within 4 days following retroviral injection. The rapid timescale of lineage reprogramming is supported by our recent transcriptomic analysis(*46*), which confirmed that most Ascl1SA6-Dlx2 iNs express markers of immature inhibitory neurons by 4 dpi (e.g. Arx, Sp9, Dcx, Gad1, Gad2, Dlx5/6). Our data also demonstrates that iNs can migrate from the earliest stages of conversion at around 2 dpi to at least 6 dpi (P11-12). The timing of morphological events also showed striking correlation with the emergence of neuron-like spontaneous calcium transients in nascent iNs at around 4 dpi (P9-10). Functional activity in endogenous cortical interneurons instructs morphological development and local migration(*33, 47*). For example, late-born CGE-derived cortical interneurons require depolarizing activity by P3 to promote migration to their correct laminar positions by P7 (*33, 47*). We speculate that Ascl1SA6-Dlx2 iNs may be undergoing a broadly similar biological process to endogenous cortical interneurons but offset by around a week.

Our retroviral vectors targeted astrocytes at a developmental time when they undergo local proliferation(*6, 7*), yet we observed only rare instances of proliferation during conversion itself. Although our data are limited to timepoints after the onset of detectable reporter protein expression (∼35-48 hpi), they nonetheless support the idea of Ascl1SA6-Dlx2 instructing rapid cell-cycle exit and neuronal differentiation, rather than a prolonged amplification stage. This did not result in a reduction of cortical astrocyte density at P17 (12 dpi). We suspect that ongoing astrogenesis allows for regulation of astrocyte density, despite reprogramming of Ascl1SA6-Dlx2 transduced astrocytes into neurons. This is important as maintenance of astrocyte numbers is almost certainly a pre-requisite to ensure long-term cortical health and function, given their essential biological roles.

Oligodendrocyte precursor cells (OPCs) proliferate in the early postnatal cortex and can be transduced by retroviral vectors(*37*) but failed to substantially contribute iNs during Ascl1SA6 mediated reprogramming in the early postnatal cortex(*19*). We cannot exclude that some neuron-like cells in the first reprogramming week are OPC-derived(*46*). However, using a robust genetic labelling approach prior to retroviral injection, we demonstrate that the vast majority, if not all, of Ascl1SA6-Dlx2 iNs were derived from the astrocytic lineage. Additionally, our transgenic approach for introducing GCaMP6s expression provides high confidence that we report the functional aspects of astrocyte-derived iNs in this work.

It has been shown that transplanted neurons from mouse embryonic cortex or derived from human stem cells can functionally integrate into mouse cortical circuits and be recruited during sensory processing(*48, 49*). Our work provides direct evidence that cortical astrocyte-derived iNs show orientation-tuned, visually evoked calcium responses to drifting gratings, suggesting they too are functionally recruited by local cortical circuitry in awake animals. Therefore, our work highlights opportunities for gene therapies to activate functional neuronal replacement in postnatal life. These could deliver useful clinical outcomes in neurodevelopmental disorders, epilepsies or perinatal cortical injuries, without the need for transplanting exogenous cellular sources or *ex vivo* cellular manipulations.

## Acknowledgments

We would like to thank Juan Burrone and Thomas Mrsic-Flogel for ideas and discussion during development and execution of this project. We are grateful to Nancy Carvajal Garcia for laboratory management and for assistance with viral production. We are also grateful to the biological services facility staff and CDN technical staff who helped to ensure animal welfare and provided necessary infrastructural support permitting this work. We thank Klaus Nave and Sandra Goebbels for their gift of the Nex-Cre transgenic line.

## Funding

MRC-ITND Studentship (MR/P502108/1; NE/W503137/1) (SL)

Wellcome Trust (206222/Z/17/Z) (AK)

European Research Council (ERC) (101021560, IMAGINE) (BB)

Wellcome Trust (206410/Z/17/Z) (BB)

German Research Foundation (BE 4182/19-1, project no. 530079744) (BB)

ERA-NET Neuron grant (Brain4Sight, 01EW2202) (BB)

Core funding to the Francis Crick Institute from Cancer Research United Kingdom, The Medical Research Council, and the Wellcome Trust (FC001002) (BB).

Epilepsy Research Institute (grant number F2404) (NM)

## Author contributions

Conceptualization: BB, AK, SL

Methodology: SL, AK, FFM, ABA, AC, NM, VS

Investigation: SL, FFP, LD, AGM, JJA, SF, NT

Visualization: SL

Funding acquisition: BB, AK

Supervision: BB, AK, NM, OM

Writing – original draft: SL

Writing – review & editing: BB, AK, with input from all authors.

## Competing interests

Authors declare that they have no competing interests.

## Data and materials availability

All data used in this work are presented in the manuscript, the Supplementary Materials, or can be downloaded from 10.18742/30920522. Plasmids/viruses and other materials generated specifically for use in this work will be provided by BB upon reasonable request. We do not report custom computation, models or simulations that go beyond common routines or analysis as described in the text or cited but code used for analysis is available on request to the lead contact(s).

## Supplementary Materials

### Materials and Methods

#### Animals

Animal procedures were approved by the King’s College London Animal Welfare and Ethical Review Body (AWERB) and conducted under the authority of the UK Home Office (Licence number: PP8849003). The following transgenic lines were used in this work: Aldh1l1-CreERT2 (IMSR_JAX:031008), RCE:loxP (MMRRC_032037-JAX), GFAP-Cre line 77.6 (IMSR_JAX:024098), Vgat-ires-Cre (IMSR_JAX:016962), Nex-Cre(*50*) and Ai96(RCL-GCaMP6s)(IMSR_JAX:028866). Reporter transgene recombination in Adlh1l1-CreERT2;RCE:loxP mice was achieved though daily administration of 75mg/kg of tamoxifen (B5965, ApexBio) dissolved in corn oil (C8267, Sigma-Aldrich) via intraperitoneal injection from P2 to P5. GFAP-Cre line 77.6 (IMSR_JAX:024098) heterozygous female animals were crossed to Ai96(RCL-GCaMP6s) homozygous male mice for *in vivo* two-photon microscopy experiments to characterise functional properties of induced neurons. C57BL6/J wild-type animals were used to visualise morphological changes during engineered neurogenesis *in vivo*.

#### Viral Vectors

Recombinant, vesicular stomatitis virus G (VSV-G)-pseudotyped, Moloney murine leukemia virus particles were prepared as previously described by transfecting modified-HEK293 retroviral packaging cell line with either pMMLV-CAG-IRES-DsRed (described in (*8*)) or pMMLV-CAG-Ascl1SA6-T2A-Dlx2-IRES-DsRed (VectorBuilder ID: VB231113-1649zjx) transfer plasmids constructed on identical viral backbones (or with identical constructs replacing DsRed with mScarlett). Briefly, packaging cell line was seeded in 10cm³ petri dishes until 80% confluent, followed by polyethylenimine-mediated (Polysciences Inc., 9002-98-6) overnight transfection. The next morning, medium was replaced (DMEM (Gibco, 11960044) supplemented with 10% FCS, 2mM L-Glutamine (Thermo Scientific, 25030024), 1mM sodium pyruvate (Gibco, 11360070), MEM Non-Essential Amino Acids (Gibco, 11140050)) and cells left at 37°C for 3 days. Viral particles were harvested by two-step ultracentrifugation of, first, filtered culture medium atop an iodixanol-based layer (OptiPrep, Sigma-Aldrich, D1556), and then re-suspended gradient layer in TBS (0.05M Tris, 0.15M NaCl, 0.01M KCl, 5mM MgCl₂) to obtain a final viral pellet. Virus was then stored at −80°C in TBS and injected at a physical titre of ∼10^8^ gc/ml.

#### Tissue preparation and immunohistochemistry

Mice were terminally anaesthetised with intraperitoneal injection of ketamine (150mg/kg) and medetomidine (20μg) and then transcardially perfused with ice-cold 0.9% sodium chloride followed immediately by 4% paraformaldehyde (PFA) in 1× phosphate-buffered saline (PBS; pH 7.4). Dissected brains were then post-fixed in 4% PFA overnight and 40 μm sections collected on a vibrating microtome (VT1000 S, Leica). Free-floating brain sections were incubated in blocking solution containing Triton™ X-100 (0.3%, v/v) and 2.5% (v/v) of donkey and goat serum (or 5% donkey serum alone) in tris-buffered saline (TBS; T6664, Sigma-Aldrich) for 1.5 hours followed by the addition of primary antibodies for overnight incubation. The following day, samples were washed with TBS and incubated for 1.5 hours in blocking solution containing secondary antibodies and then mounted onto microscopy slides with Mowiol® (81381, Sigma-Aldrich) prior to imaging.

The following antibodies were used in this work: Chicken Anti-GFP Polyclonal (Antibodies Incorporated Cat No GFP-1020, RRID:AB_10000240), Rabbit Anti-RFP pre-adsorbed (Rockland Cat No 600-401-379, RRID:AB_2209751), Mouse Anti-Sox9 Monoclonal (Thermo Fisher Scientific Cat No 14-9765-80, RRID:AB_2573005), Mouse Anti-NeuN (Sigma-Aldrich Cat No MAB377, RRID:AB_2298772), Alexa Fluor™ 488 AffiniPure Donkey Anti-Chicken IgY (IgG) (H+L) (Jackson ImmunoResearch Labs Cat No 703-545-155, RRID:AB_2340375), Goat anti-Rabbit IgG (H+L) Cross-Adsorbed Secondary Antibody, Alexa Fluor™ 568 (Thermo Fisher Scientific Cat No A-11011, RRID:AB_143157) and Goat anti-Mouse IgG1 Cross-Adsorbed Secondary Antibody, Alexa Fluor™ 647 (Thermo Fisher Scientific Cat No A-21240, RRID:AB_2535809).

#### Animal Surgeries

Neonatal (P5-6) mice were anaesthetised using 5% isoflurane (1L/min) for 2–3 minutes for induction and 1–2% for maintenance throughout surgical procedures with body temperature maintained at 37°C. Animals were placed in a stereotaxic apparatus using rubber head bars. Around 30 minutes prior to induction, topical 5% lidocaine ointment was applied to the scalp over the surgical site. Aseptic technique and regular assessment of anaesthetic depth were employed throughout surgeries.

For stereotactic cortical injections, a minimal (1–2mm) sagittal scalp incision was made approximately 1mm right of midline overlying posterior cortical regions. A small nick in the skull was made with a sterile needle at +2mm (ML) from lambda to allow passage of the injection micropipette. 1μL of viral suspension was injected using a pulled glass micropipette over 10 minutes at 0.3-0.4mm from the cortical surface and then slowly retracted over several minutes. Skin was then closed and secured with tissue adhesive (Vetbond™, 3M™) and the mouse allowed to recover before returning to home cage. For neonatal cranial window and headplate implantation, a region of scalp skin was removed to expose the skull surface above the intended implantation site. A custom-made headplate containing a 3.5mm diameter recess was fixed to the skull over the underlying visual cortex using cyanoacrylate adhesive (Loctite) and surrounding skin secured. A craniotomy of 3mm diameter was performed to expose the posterior cortical surface. 1μL of viral suspension was injected using a pulled glass micropipette over 10 minutes at ∼2mm ML from lambda and 0.3-0.4mm from the cortical surface and then slowly retracted over several minutes. A small drop of Kwik-Sil™ ( WPI) was placed at the centre of the site and immediately covered with a 3mm diameter, 0.1mm thick glass coverslip (64-0720, Warner Instruments). The perimeter of the glass was secured, using cyanoacrylate adhesive, to the surrounding skull and headplate. Finally, the headplate was further secured to the skull using dental cement (Super-Bond C&B Kit, Sun Medical). Buprenorphine (0.05mg/kg, subcutaneous) was administered for post-operative analgesia and monitored over the subsequent 48 hours for post-operative complications.

Juvenile mice (>P12) cranial window and headplate implantation proceeded largely similarly to neonates with the exception that a prior intraperitoneal injection of meloxicam (5mg/kg) was administered for intraoperative analgesia and Viscotears® Liquid Gel eye drops were applied to both eyes during the procedure. Juvenile mice did not receive concurrent cortical injection of virus, rather the window was placed over the visual cortex having previously received viral injection at the site at postnatal day 5-6.

#### Confocal Imaging and Quantification of Fixed Tissue

Microscopy data were collected on Zeiss Axio Imager.Z2 LSM 800 confocal microscope equipped with four solid-state lasers (405, 488, 561, and 633 nm) using a Plan-Apochromat 20x/0.8 air objective (Zeiss, 440640-9903-000) and z-spacing of 1 μm. Direct comparison (images and quantification) between groups were conducted on images collected with identical microscopy parameters. Manual cell counts were performed in ImageJ. Blinding of counts during analysis was not practicable due to strikingly obvious differences between the morphology of RFP+ cells across groups. Estimates of cell density were conducted on images acquired through entire z-axis profile of fixed tissue by first manually defining an ROI at the injection site, then segmenting Sox9 images using Cellpose(*51*) at this ROI and in the contralateral hemisphere in a comparable anatomic region of cortex. The number of segmented Sox9+ cell soma were then divide by the tissue volume, calculated by taking the 2D image/ROI area in microns and multiplying this by the wet tissue thickness depth of 40 μm.

#### Explant Culture and Time-lapse confocal microscopy

Postnatal brains were dissected in ice-cold 1X Krebs buffer (NaCl 0.125M, KCl 2.5mM, NaH2PO4 1.2mM, MgCl2 1.2mM, CaCl2 2.5mM), embedded in 4% agarose and sectioned to a thickness of 300 μm on a vibratome (Leica VT1000S) in ice-cold oxygenated (95%) 1X Krebs buffer. Coronal brain slices were placed in a culture filter membrane insert (Millipore) inside a glass-bottom culture dish (FluoroDish, World Precision Instruments) containing serum-supplemented medium consisted of DMEM with 10% FBS, 45% Glucose and Pen-Strep (100 U ml−1). Slices were incubated during 1h at 37C, 5% CO2. After incubation, medium was replaced by serum-free medium consisting of Neurobasal with B27 supplement (1:50; Invitrogen), N2 Supplement (1:100; Invitrogen), 45% Glucose, GlutaMax (1:100; Invitrogen), Pen-Strep (100 U ml−1) and place for imaging on a Zeiss LSM880 confocal microscope equipped with stage-top incubator at 37 °C with a 5% CO2 atmosphere. Brain slices were imaged using a 10x objective with a time interval of 20 min between images.

#### Two-photon Microscopy Acquisition and Data Pre-processing

Two-photon structural imaging and spontaneous calcium imaging was collected on a custom-built resonant scanning two-photon microscope (Cosys) operated by Scanimage (2023.0.0, MBF Bioscience) using Chameleon Discovery NX tunable laser (Coherent) at 920 nm wavelength for GCaMP6s functional imaging and 1060 nm wavelength for structural image acquisition. Images were collected using a 25X (NA 1.05, OFN 18) water-immersion objective (Olympus, XLPLN25XWMP2). Functional images were acquired at 512×512 pixel resolution at a framerate of 30Hz. Ascl1SA6-Dlx2 iNs were identified prior to functional recordings by imaging at 1060 nm to corroborate RFP expression and iN morphology. Structural images were collected at 1024×1024 resolution with 250 frames acquired per slice. To follow conversion of superficial cortical glia into neurons, stacks were acquired from below the pial surface down to approximately 200 μm. Blood vessel anatomy was used to re-identify imaging sites on subsequent days. Mice were head fixed under the microscope objective on a stable platform but with freedom to move their trunk, limbs, tail etc. Neonatal mice (<P12) were placed over a heating pad (32°C) throughout imaging sessions.

For assessing responses to visual stimuli, images were recorded on custom-built resonant scanning two-photon microscopes (Cosys) with either a Chamelean Vision S laser and 12 kHz resonant scanners (Cambridge Technology) or a Mai Tai High-Performance laser and 9 kHz resonant scanners (Cambridge Technology), both operated by Scanimage (MBF Bioscience) using an FPGA module (PXIe-7965R FlexRIO, National Instruments). Functional images were recorded at 920 nm using a 16X, 0.8NA objective (Nikon) and the FOV size adjusted for each ROI or groups of ROIs. Visual stimuli were generated using Psychtoolbox-3 in MATLAB. Mice were head-fixed and placed on either a platform or a running wheel and were shown sinusoidal visual gratings drifting in one of eight directions (incremental rotations of 45°), with a spatial frequency of 0.1 cycles per degree and a temporal frequency of 2Hz. An additional blank (no gratings) stimulus was also included. The total of nine visual stimuli conditions were then randomly presented for a duration of 2 seconds with a 5 second interval before the start of the next visual stimulus presentation. An average of 10 trials per stimulus was shown.

Two-photon imaging data pre-processing was performed in Python using Suite2p(*52*) for image registration (motion correction) within planes. For structural volumetric images to track engineered neurogenesis in the cortex (**Fig. 1**), we used StackReg(*53*) plugin in FIJI, ImageJ(*54*) to align planes prior to downstream analysis. For functional imaging analyses, peri-somatic cellular ROIs were manually constructed in Labkit(*55*) based on the mean fluorescence signal from green (GCaMP6s), using red channel (RFP) fluorescence and morphology to distinguish neighbouring astrocytes from retroviral transduced cells expressing RFP reporter protein. Following ROI construction, signal (raw fluorescence (Fr) and neuro-/glial-pil (Fneu)) extraction was performed in Suite2p(*52*). Contaminating neuro-pil/glial-pil signal was removed by subtraction of 0.7×Fneu from Fr. For spontaneous calcium activity analysis, a moving baseline fluorescence (F₀) was calculated using Suite2p’s minimax implementation with default parameters (suite2p v.0.14.3) and ΔF/F was calculated as (F – F₀) / F. For calcium imaging in the early neonatal period (<7 dpi) after neuropil correction, a constant mean of each ROIs neuropil trace was added back to the corrected F signal as a pragmatic measure to avoid spurious ΔF/F spikes caused by division by near-zero values in cells with low activity and low baseline fluorescence. For orientation mapping, F₀ was calculated by taking the 2.5th percentile of the glia-/neuro-pil corrected fluorescence signal after median filtering with a 0.75-second window in MATLAB. ΔF/F was then calculated as (F – F₀) / F₀.

### Data Analysis and Statistics

#### Morphology and migration

Following registration and stack alignment, we used semi-automated rooted tree-structure reconstructions of RFP+ transduced cells were performed using Simple Neurite Tracer(*7*) in ImageJ. Measurements of branch length, primary branch number, longest-shortest path, number of tips (endpoints within the tree structure) as described in SNT documentation(*7*). We performed separate regression models for each outcome variables association with time ( hpi) using statsmodels(*56*) (v0.14.5) in Python. Continuous outcomes were standardized (z-scored) prior to analysis and modelled using ordinary least squares (OLS) regression (*zY*_*i*_ = *β*_0_ + *β*_1_ × (ℎ*pi*_*i*_) + *ε*_*i*_). For discrete count variables (e.g., number primary branches) we used Poisson regression with a log link function (*Y*_*i*_ ∼ *Poisson*(*μ*_*i*_); *log*(*μ*_*i*_) = *β*_0_ + *β*_1_ × (ℎ*pi*_*i*_)). In both cases employing heteroskedasticity-robust standard errors(*9*). Results, including beta coefficients and Benjamini–Hochberg false discovery rate (FDR)(*10*) corrected p-values (α = 0.05) are presented in **Table S1**.

Quantification of migration speed and duration was performed manually in 2D using maximum intensity projections of coronal sections acquired using time-lapse confocal microscopy. We selected individual cells that moved locations during playback and measured the full distance of the path of the cell soma taken using ImageJ from their initial location until the cell ceased to migrate further. The minimum time interval taken for a cell to move from its starting location to a stationary destination (or out of frame/end of recording) was used to define the duration of a migration event and used to calculate migration speed. Thus, measures of migration speed and distances in the text represent the averaged speed of actual migration, excluding stationary periods.

#### Analysis of spontaneous calcium activity

To define spontaneous calcium events (i.e. events in the absence of experimentally controlled stimulus presentation or behavioural paradigms; **Fig. 2, 4**), we used a two-state Gaussian hidden Markov model (HMM) with predefined, fixed parameters to identify ‘active’ states in which the signal exceeded 2.5 standard deviations above baseline. ΔF/F traces were z-scored and smoothed using a Savitzky–Golay filter. The model comprised a baseline state (mean = 0) and an active event state (mean = 2.5, corresponding to 2.5 SD above the mean of the z-scored ΔF/F signal), with diagonal covariance (σ² = 0.1) for both states. State transitions were constrained by high self-transition probabilities (0.9) and low switching probabilities (0.1), enforcing temporal persistence. The most likely sequence of hidden states was inferred using the Viterbi algorithm and implemented in hmmlearn (v0.3.3, Python). ROIs that had zero calcium events were excluded from subsequent analyses measuring or comparing properties of such events.

To statistically assess changes in calcium event frequency over time (mean centered) in the early stages of lineage reprogramming (**Fig. 2**), we fitted an ordinary least squares (OLS) linear regression model with robust standard errors(*9*) (**Table S2**) in statsmodels:

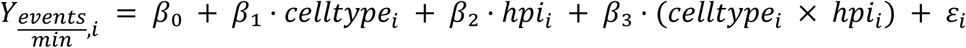

Where:

- β0 (intercept): mean event frequency for astrocytes at mean timepoint ( hpi),
- β1 baseline difference between cell types (non-transduced astrocytes vs. Ascl1SA6-Dlx2 transduced cells),
- β2 linear effect of time for astrocytes,
- β3 interaction coefficient between time and cell type.

To statistically assess mean differences in Pearson’s correlations across multiple categorical groups of cell type pairs, we performed a one-way ANOVA followed by pairwise Tukey’s test in statsmodels. To estimate confidence intervals of between-group, unpaired mean differences in calcium event frequency, duration and amplitudes we performed bootstrap resampling (5000 iterations, bias-corrected and accelerated intervals) using dabest(*57*).

#### Analysis of visually evoked calcium activity

Glia-/neuro-pil corrected ΔF/F for constructed ROIs were aligned to stimulus onset. For each stimulus presentation trial, we defined a baseline pre-stimulus window as the one second interval preceding the stimulus onset. The mean and standard deviation (SD) of activity in this window were used to z-score the post-stimulus responses (which we defined as the 2.5 seconds following stimulus onset). We averaged trials where the same stimulus (i.e. drifting grating orientation or blank stimulus) was shown to get mean responses to a given stimulus. We excluded all recordings where the mean number of trials per stimulus was <8 from downstream analysis. We calculated response magnitudes to a given stimulus as the difference in the mean of the trial-averaged z-scored activity of pre- and post-stimulus periods. Using the response magnitudes of the blank (grey screen, no grating) stimulus condition for every cell in the entire dataset as a control reference distribution (i.e. distribution of response magnitudes in the absence of drifting gratings) we computed a one-sample t-statistic and p-value for each individual cell’s response magnitude to all 8 orientations of the drifting grating stimuli (*t*_*c*_ = (*Δ*_*c*_ − *μ*_*blank*_) / (*s*_*blank*_ ∗ √(1 + 1/*n*_*blank*_)), where Δ_c_ is the response magnitude for a given cell and stimulus and μ, s and n are the mean, SD and number of cells in the blank reference distribution). We defined a cell as “responsive” to a given stimulus if the p-value corresponding to the t-statistic was lower than a Bonferroni adjusted significance threshold (adjusted for the number of stimuli (i.e. 8) and the number of cells being tested; α_Bonferroni_ = 0.05/8*N_cells_). The preferred stimulus of a cell was defined as the orientation at which that cell showed the greatest positive response magnitude. Orientation selectivity index (OSI) of a cell was calculated as:

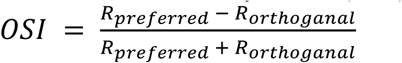

where R_preferred_ was the response magnitude at the preferred orientation and R_orthoganal_ was the mean of the response magnitudes at orientations at 90 degrees (in either direction) to the preferred stimulus. To determine whether cell types showed greater response magnitudes to the stimulus in the opposite direction (i.e. 180 degrees) to the preferred stimulus than other orientations, we calculated the difference between the opposite orientation at 180 degrees (R_opposite_) and the mean magnitude of the responses to all other orientations (excluding R_preferred_) for each cell. This elicited a contrast value for each cell, which we could then use to compute a one-sample, one-tailed t-test to test the null hypothesis that the contrast was zero (i.e. no preference in response of the opposite (180 degrees of rotation) grating compared to the remaining orientations (**Table S3**).

#### Statistical reporting

Means and 95% confidence intervals for nested proportions/percentages (e.g. percentage of cells expressing a marker protein that are clustered within individual mice) were calculated on the logit scale and back-transformed (inverse logit) to original percentage scale for reporting. Estimates of mean differences between groups from quantifications made from histological data were performed in dabest using bootstrap resampling (5000 iterations, bias-corrected and accelerated intervals) to obtain 95% confidence intervals. Two-sided permutation testing (with 5000 reshuffles of group labels) was used to obtain the probability (p) of observing at least the magnitude of difference if the null hypothesis of no difference between groups were true.

Unless otherwise stated in legends, graphs visualizing regression lines in the Figures used seaborn’s regplot with default parameters where regression lines calculated by ordinary least square regression with shaded regions depicting 95% confidence intervals estimated from 1000 bootstrapped resamples. Categorical boxplots whiskers are drawn up to 1.5 x IQR (interquartile range) from the nearest box hinge (marking either 25^th^ or 75^th^ percentile of the data) or until the minimum or maximum range of the data if it is within this interval. Wherever p-values have been reported in this work following hypothesis testing, α was defined at 0.05 (unless specifically stated otherwise).

**Fig. S1.**
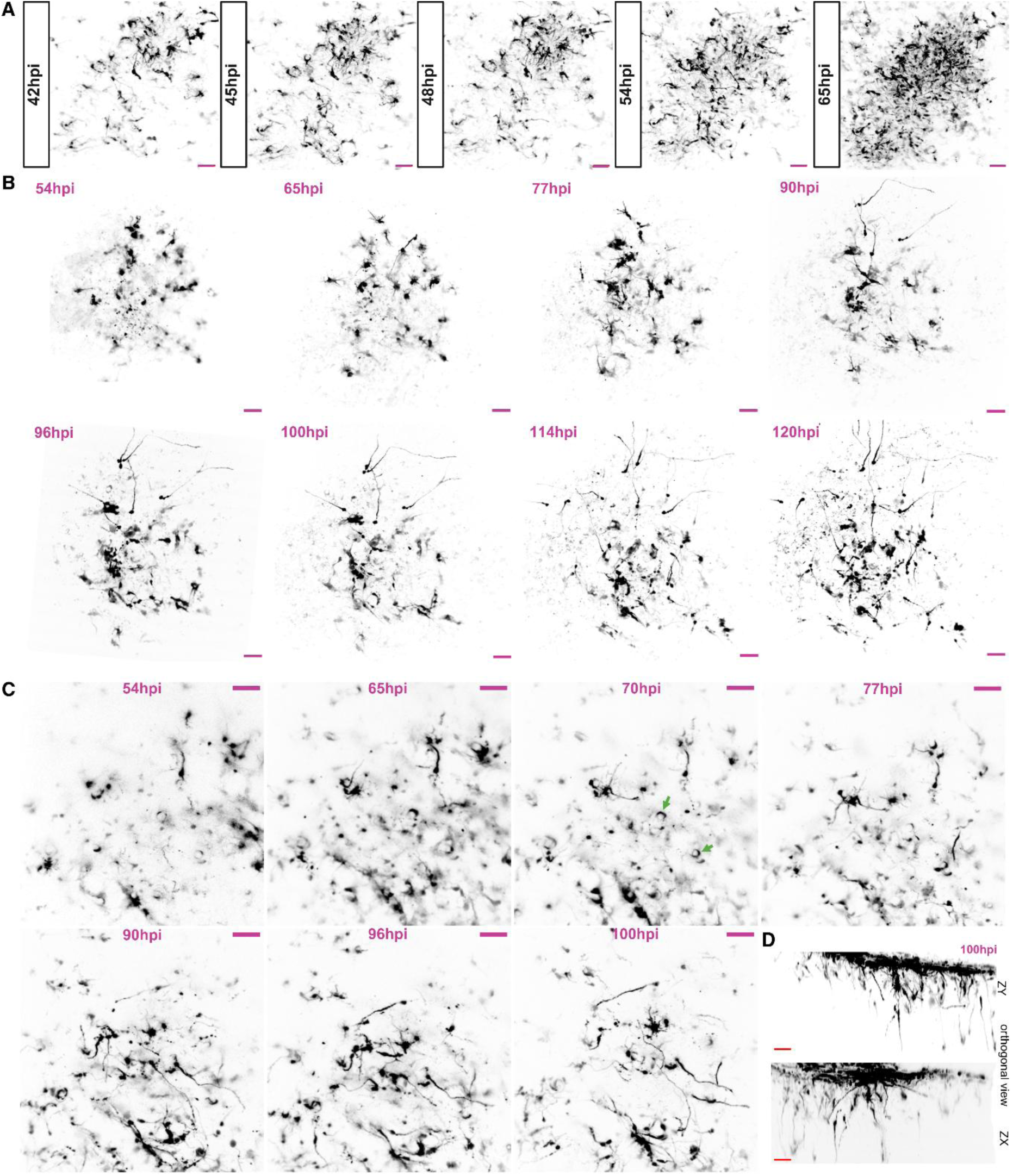
Longitudinal 2-photon imaging of glia to neuron reprogramming with Ascl1SA6-Dlx2. **(A)** Expression of RFP from MMLV-Ascl1SA6-T2A-Dlx2-IRES-RFP can be detected from around 40 hpi, gradually increasing over the course of the subsequent ∼24 hours. **(B)** Full time series of max. intensity projections (MIP) of image volumes, partially presented in Fig.1D**. (C)** Additional example of MIP of image stacks from 54-100 hpi in a third animal. Green arrows in panel at 70 hpi indicate examples of RFP+ glial cells ensheathing blood vessels. **(D)** Orthogonal MIP views of image volume acquired at 100 hpi.

**Fig. S2.**
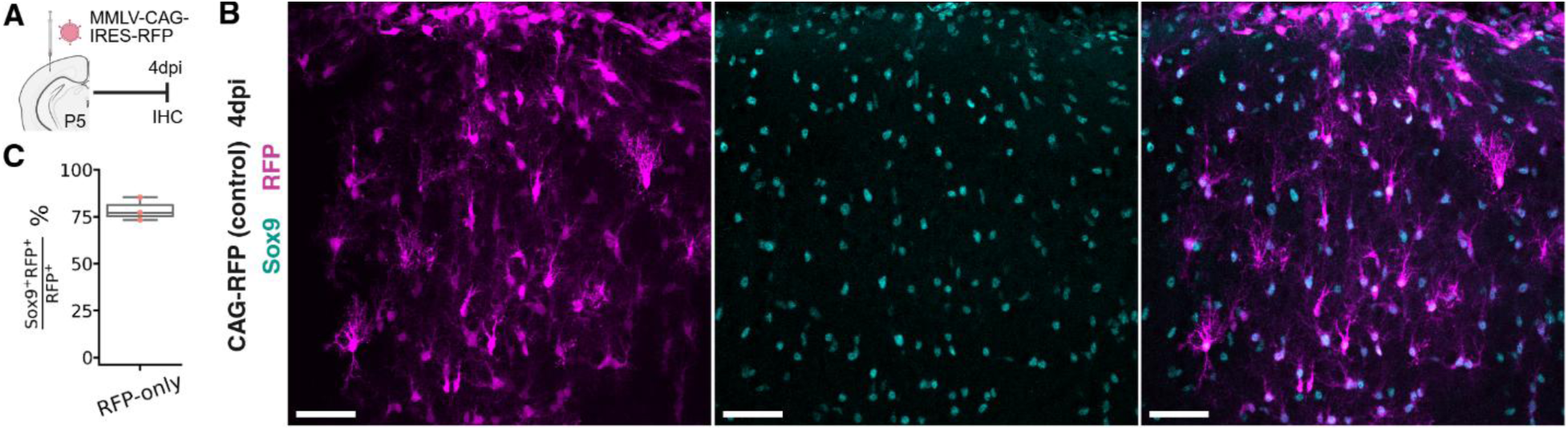
RFP-only control virus transduced cells are glia and predominantly Sox9+ 4 dpi. **(A)** Schematic. MMLV-CAG-IRES-RFP was injected into wild-type C56BL/J P5-6 pups with analysis on tissue fixed at 4 dpi. **(B)** Example of the morphologies of RFP+ cells in stained coronal sections of mouse cortex (magenta) and Sox9 staining (cyan). **(C)** Percentage of RFP+ cells in MMLV-CAG-IRES-RFP (control) injected cortices that express Sox9 (mean 79.04%, 95%CI [70.86%, 85.39%]).

**Fig. S3.**
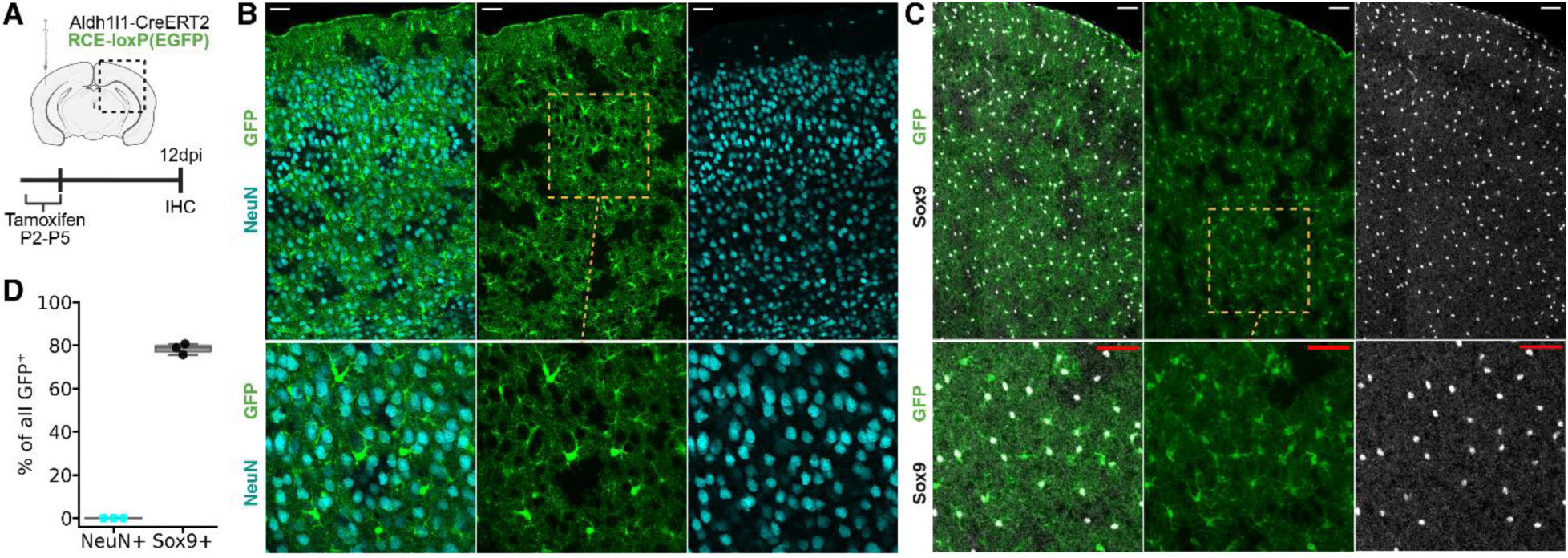
Aldh1l1-CreERT2 restricts Cre-dependent GFP (RCE:loxP) expression to astrocytes in neonatal mouse cortex. **(A)** Experimental design. **(B)** Single plane of confocal image of coronal slice of hemisphere contralateral to injection of virus stained for expression of NeuN (cyan) and GFP (green). **(C)** MIP of confocal stack of coronal slice of hemisphere contralateral to injection of virus stained for expression stained for Sox9 (gray). **(D)** Quantification of the proportion of all GFP+ cells that co-express NeuN (cyan) or Sox9 (black/gray). N=3 mice, >4500 cells. No examples of GFP+NeuN+ cells. Sox9 expressed in 78.48% (95%CI [75.50%, 81.19%]) of GFP+ cells. Scale bars: 50 μm.

**Fig S4.**
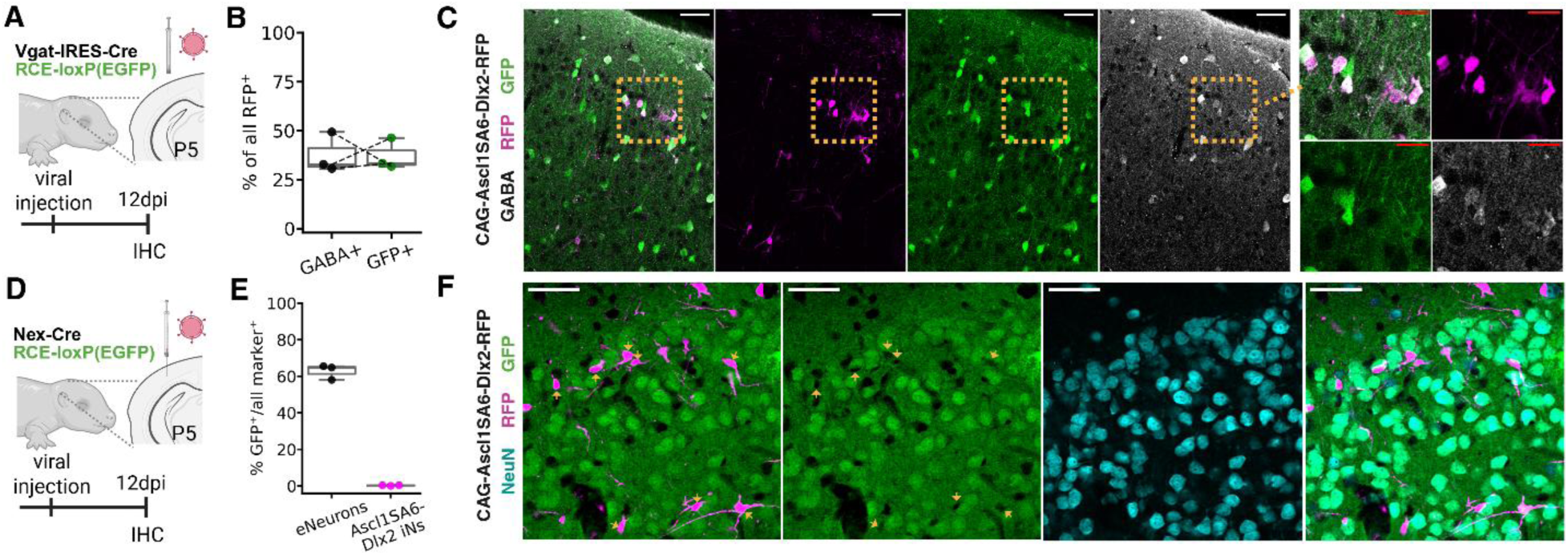
Ascl1SA6-Dlx2 iNs co-localise with labels of inhibitory, but not glutamatergic, neuronal identity. **(A)** Experimental design for pan-GABAergic neuronal labelling. **(B)** Quantification of the percentage of RFP+ Ascl1SA6-Dlx2 transduced cells that co-express GABA and GFP in Vgat-ires-Cre;RCE:loxP(GFP) mice. **(C)** MIP showing an example section of cortex with RFP+ Ascl1SA6-Dlx2 iNs co-stained for GFP and GABA. **(D)** Experimental design for pan-glutamatergic neuronal labelling. **(E)** Quantification of the percentage of endogenous neurons (eNeurons; black dots – i.e NeuN+ neurons that were not magenta/Ascl1SA6-Dlx2 iNs) and RFP+ Ascl1SA6-Dlx2 iNs (magenta dots) that co-express GFP in Nex-Cre;RCE:loxP(GFP) mice. **(F)** MIP showing an example section of cortex used for quantification in (E) with Ascl1SA6-Dlx2 RFP+ cells co-stained for NeuN and (absent) GFP (yellow arrowheads).

**Fig. S5.**
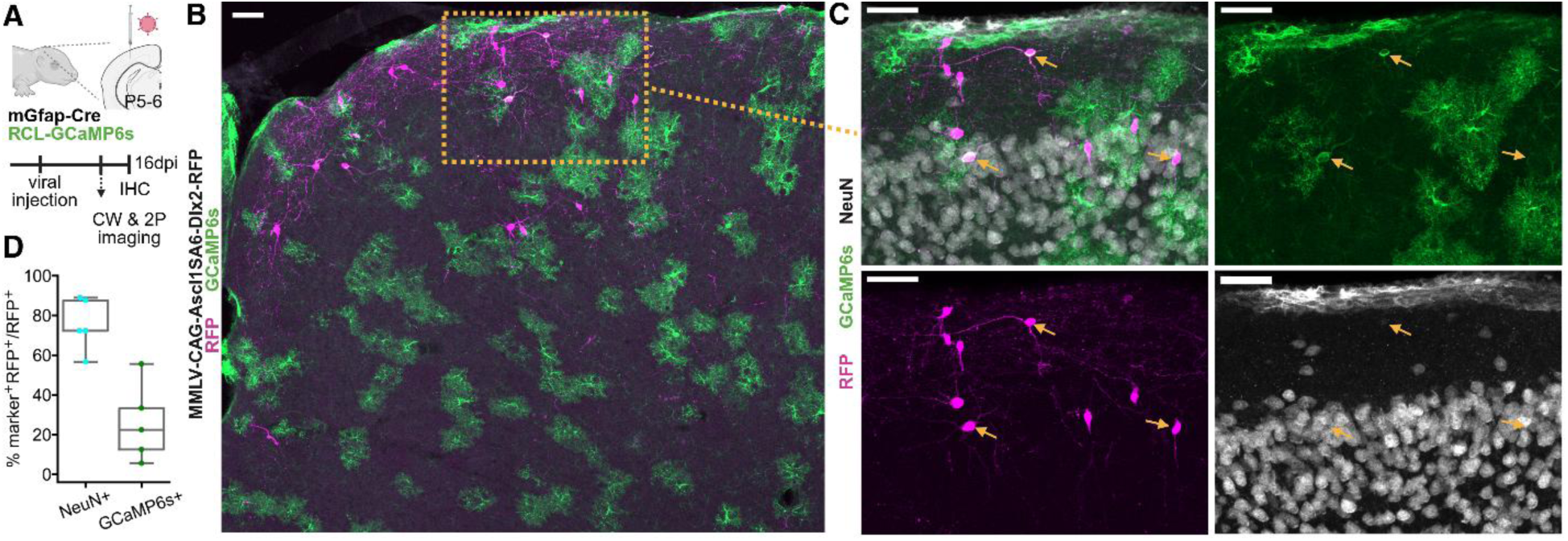
Ascl1SA6-Dlx2 iNs express cre-dependent GCaMP6s in mGfap-Cre;Ai96 mice. **(A)** Experimental design. **(B)** MIP of injected hemisphere from a representative animal used for quantification in (D). **(C)** MIP of magnified section indicated from (B) showing RFP (magenta), GCaMP6s (green) and NeuN (gray) expression. **(D)** Quantification of the proportion of RFP+ cells that express NeuN (cyan; (77.56%, 95%CI [63.95%, 87.07%])) and the proportion of Ascl1SA6-Dlx2 iNs (RFP+NeuN+ cells) that express GCaMP6s (green, (21.44%, 95%CI [8.93%, 43.18%])). N=4 mice, 202 RFP+ cells counted.

**Fig. S6.**
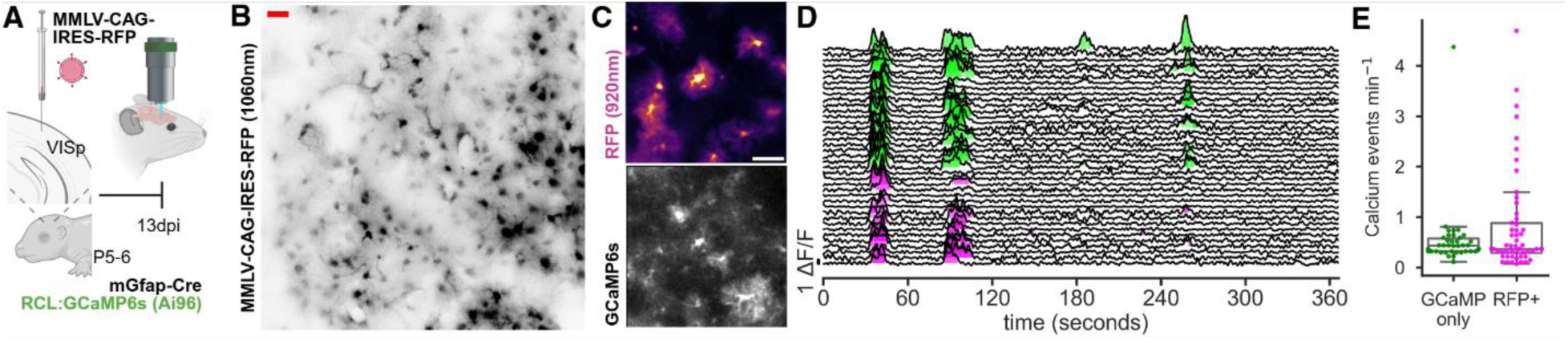
RFP-only control astrocytes show glial morphology and low frequency, astrocyte-like calcium dynamics at 13 dpi. **(A)** Experimental schematic. **(B)** MIP of a 2-photon structural stack acquired at 1060 nm showing glial-like morphologies at 13 dpi. **(C)** Example mean intensity projection images of RFP and GCaMP6s expression from a spontaneous imaging recording session recorded at 920 nm following RFP-only control retroviral transduction at 13 dpi. **(D)** Representative spontaneous calcium activity recorded from RFP-only transduced astrocytes (magenta events shaded) and neighboring GCaMP6s+ astrocytes (green events shaded). **(E)** Calcium event frequency in RFP-only control transduced astrocytes and neighboring astrocytes from the same recordings at 13 dpi.

**Table S1.** Morphological dynamics over time post-injection of Ascl1SA6-Dlx2 retrovirus. Model results estimating relationships between time ( hpi) and listed outcome variable. Continuous outcome variables where OLS regression was implemented were z-cored prior to model fitting, thus beta coefficients can be interpreted as change in standard deviations of the outcome variable for an additional hour of time post-injection. Beta coefficients for the Poission GLM (log link) should be interpreted as the log rate ratio (i.e. the exponent of the coefficient multiplicative change in the expected count for a one hour increase in time post-injection; e.g. exp(**β_1_**)=1 indicates no change, exp(**β_1_**)=0.5 indicates halving of expected count over an hour). We report R² for OLS regression and Cox–Snell pseudo-R² for Poisson GLM for models of count outcomes.

| Variable | Model | Beta ($\beta_1$) | 95% CI | SE | Z | p-value (FDR-corrected) | $R^2$ / ↑pseudo$R^2$ |
| --- | --- | --- | --- | --- | --- | --- | --- |
| No. of primary branches | Poisson GLM | -0.00657 | [-0.00851,-0.00464] | 0.000987 | -6.65 | 3.47E-11 | 0.07<sup>†</sup> |
| No. of tips | Poisson GLM | -0.01072 | [-0.01342,-0.00802] | 0.001377 | -7.78 | 1.16E-14 | 0.27<sup>†</sup> |
| Mean branch length (z-scored, SD 43.7 $\mu\text{m}$) | OLS | 0.017362 | [0.013664,0.021061] | 0.001887 | 9.20 | 8.91E-20 | 0.17 |
| Length of longest shortest path (z-scored, SD 63.7 $\mu\text{m}$) | OLS | 0.017318 | [0.013815,0.020821] | 0.001787 | 9.69 | 1.66E-21 | 0.17 |
| Convex hull: Size (z-scored, SD $5.3 \times 10^4 \mu\text{m}^3$) | OLS | 0.006167 | [0.001722,0.010612] | 0.002268 | 2.72 | 0.006539 | 0.02 |

**Table S2.**
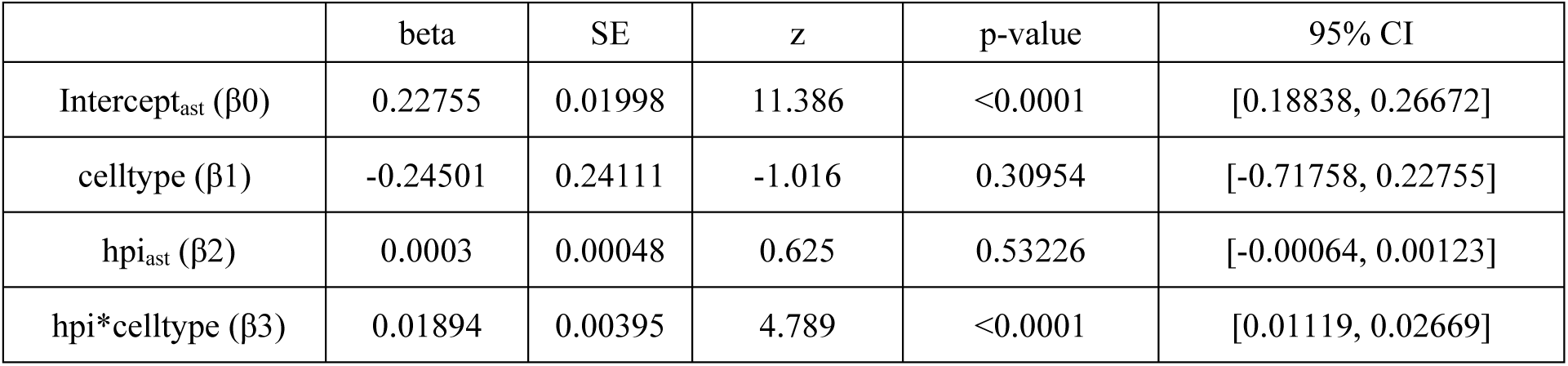
Results from linear regression that was used to test whether the interaction between time post-injection ( hpi – variable centered such that intercept is at 48 hpi) and cell type (Ascl1SA6-Dlx2 transduced (RFP+GCaMP6s+) vs non-transduced astrocytes (RFP-GCaMP6s+)) significantly predicted calcium event frequency (see **Fig. 2G** and materials and methods). Intercept parameter β0 indicates the model prediction of the mean calcium event frequency (events min^-1^) for non-transduced astrocytes at 48 hpi. β1 is the estimate for the difference in calcium event frequency between Ascl1SA6-Dlx2 transduced and non-transduced astrocytes (implies 0.2275 – 0.2450 ≈ – 0.0175 events min^-1^ at 48 hpi for Ascl1SA6-Dlx2 transduced cells, suggesting near absent activity, although not significantly different from local non-transduced astrocytes, p=0.31). β2 parameter is the slope of the relationship between time and event frequency for astrocytes (near 0). β3 estimates the additional slope for Ascl1SA6-Dlx2 transduced cells compared to non-transduced astrocytes (indicating 0.0003+0.01894= 0.01924 events min^-1^ increase per hour, or ∼0.46 events min^-1^ per day). Overall, the regression was statistically significant F=13.10 (p<0.001), adj. R^2^ = 0.491. N_cells_ = 217 (after exclusion of 81 ROIs with 0 calcium events), dof = 3. (α = 0.05).

**Table S3.**
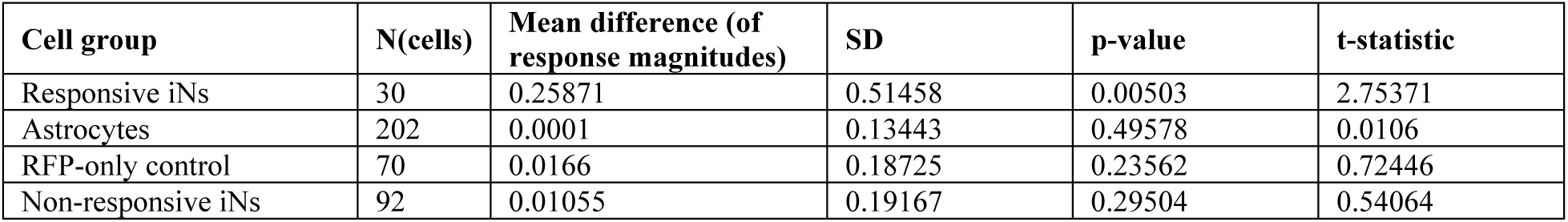
Results from one-sample t-test to determine the probability of observing the contrast between the response magnitude at the opposite direction (i.e. 180 degree orientation) to the preferred orientation and the mean response magnitude to the other orientations (excluding the preferred orientation) if the true contrast was zero under the null hypothesis (test was one-tailed (greater response at 180 degrees; alpha = 0.05/4 cell groups). (α = 0.05).

**Movie S1.**

Example of time-lapse confocal imaging from explant cortical slice in culture previously transduced with Ascl1SA6-Dlx2-RFP at P5. Data presented between 2 and 3 dpi (45-72 hpi) with several representative examples of migrating Ascl1SA6-Dlx2 iNs during lineage conversion. Even within this limited time frame, many cells also show general phenomena described from data acquired *in vivo* (**Fig. 1**) of dynamic remodeling of shorter hair-like processes with gradual neurite extension (typically one or few neurites)

**Movie S2.**

Related to Figure 2. Example of Ascl1SA6-Dlx2 iN at 6 dpi (150 hpi) and surrounding GCaMP6s+ astrocytes. Recording playback at 10x speed. Mean filter applied over 1s.

**Movie S3.**

Movies play through imaging planes (from superficial to deep) from structural imaging stacks acquired at 1060 nm following injection of retrovirus encoding Ascl1SA6-Dlx2-RFP. Two examples from 2 different mice at 4-5 hpi (100 hpi and 120 hpi, respectively), followed by two examples from 2 different mice at 13 dpi.

**Movie S4.**

Example of spontaneous GCaMP6s fluorescence recording acquired at 13 dpi in an Ascl1SA6-Dlx2 iN and nearby GCaMP6s+ astrocytes. Recording playback at 10x speed. Mean filter over 300ms.

**Movie S5.**

Example of GCaMP6s fluorescence recorded in Ascl1SA6-Dlx2 iNs at 2 wpi, illustrating visually evoked calcium transients. Inset graphics depict stimulus presentation onset, offset and orientation of drifting gratings (or blank). Recording playback at 8x speed. Mean filter applied over 500ms. Magenta arrows indicate RFP+ visually responsive induced neuron.

## Notes

### Competing Interest Statement

The authors have declared no competing interest.

## References

1. B. J. Molyneaux, P. Arlotta, J. R. Menezes, J. D. Macklis, Neuronal subtype specification in the cerebral cortex. Nat Rev Neurosci 8, 427–437 (2007).

2. O. Marín, Development of GABAergic Interneurons in the Human Cerebral Cortex. Eur J Neurosci 61, e70136 (2025).

3. S. C. Noctor, A. C. Flint, T. A. Weissman, W. S. Wong, B. K. Clinton, A. R. Kriegstein, Dividing precursor cells of the embryonic cortical ventricular zone have morphological and molecular characteristics of radial glia. J Neurosci 22, 3161–3173 (2002).

4. T. E. Anthony, C. Klein, G. Fishell, N. Heintz, Radial glia serve as neuronal progenitors in all regions of the central nervous system. Neuron 41, 881–890 (2004).

5. P. Malatesta, E. Hartfuss, M. Gotz, Isolation of radial glial cells by fluorescent-activated cell sorting reveals a neuronal lineage. Development 127, 5253–5263 (2000).

6. W. P. Ge, A. Miyawaki, F. H. Gage, Y. N. Jan, L. Y. Jan, Local generation of glia is a major astrocyte source in postnatal cortex. Nature 484, 376–380 (2012).

7. S. Clavreul, L. Abdeladim, E. Hernandez-Garzon, D. Niculescu, J. Durand, S. H. Ieng, R. Barry, G. Bonvento, E. Beaurepaire, J. Livet, K. Loulier, Cortical astrocytes develop in a plastic manner at both clonal and cellular levels. Nat Commun 10, 4884 (2019).

8. C. Heinrich, R. Blum, S. Gascon, G. Masserdotti, P. Tripathi, R. Sanchez, S. Tiedt, T. Schroeder, M. Gotz, B. Berninger, Directing astroglia from the cerebral cortex into subtype specific functional neurons. PLoS Biol 8, e1000373 (2010).

9. B. Berninger, M. R. Costa, U. Koch, T. Schroeder, B. Sutor, B. Grothe, M. Gotz, Functional properties of neurons derived from in vitro reprogrammed postnatal astroglia. J Neurosci 27, 8654–8664 (2007).

10. N. Heins, P. Malatesta, F. Cecconi, M. Nakafuku, K. L. Tucker, M. A. Hack, P. Chapouton, Y. A. Barde, M. Gotz, Glial cells generate neurons: the role of the transcription factor Pax6. Nat Neurosci 5, 308–315 (2002).

11. S. Sirko, C. Schichor, P. Della Vecchia, F. Metzger, G. Sonsalla, T. Simon, M. Burkle, S. Kalpazidou, J. Ninkovic, G. Masserdotti, J. F. Sauniere, V. Iacobelli, S. Iacobelli, C. Delbridge, S. M. Hauck, J. C. Tonn, M. Gotz, Injury-specific factors in the cerebrospinal fluid regulate astrocyte plasticity in the human brain. Nat Med 29, 3149–3161 (2023).

12. D. Hampton, K. Rhodes, C. Zhao, R. Franklin, J. Fawcett, The responses of oligodendrocyte precursor cells, astrocytes and microglia to a cortical stab injury, in the brain. Neuroscience 127, 813–820 (2004).

13. S. Bardehle, M. Kruger, F. Buggenthin, J. Schwausch, J. Ninkovic, H. Clevers, H. J. Snippert, F. J. Theis, M. Meyer-Luehmann, I. Bechmann, L. Dimou, M. Gotz, Live imaging of astrocyte responses to acute injury reveals selective juxtavascular proliferation. Nat Neurosci 16, 580–586 (2013).

14. A. Buffo, M. R. Vosko, D. Erturk, G. F. Hamann, M. Jucker, D. Rowitch, M. Gotz, Expression pattern of the transcription factor Olig2 in response to brain injuries: implications for neuronal repair. Proc Natl Acad Sci U S A 102, 18183–18188 (2005).

15. S. Gascon, E. Murenu, G. Masserdotti, F. Ortega, G. L. Russo, D. Petrik, A. Deshpande, C. Heinrich, M. Karow, S. P. Robertson, T. Schroeder, J. Beckers, M. Irmler, C. Berndt, J. P. Angeli, M. Conrad, B. Berninger, M. Gotz, Identification and Successful Negotiation of a Metabolic Checkpoint in Direct Neuronal Reprogramming. Cell Stem Cell 18, 396–409 (2016).

16. Z. Guo, L. Zhang, Z. Wu, Y. Chen, F. Wang, G. Chen, In vivo direct reprogramming of reactive glial cells into functional neurons after brain injury and in an Alzheimer’s disease model. Cell Stem Cell 14, 188–202 (2014).

17. C. Heinrich, M. Bergami, S. Gascon, A. Lepier, F. Vigano, L. Dimou, B. Sutor, B. Berninger, M. Gotz, Sox2-mediated conversion of NG2 glia into induced neurons in the injured adult cerebral cortex. Stem Cell Reports 3, 1000–1014 (2014).

18. F. Chen, X. Liu, X. Zhong, X. Chen, E. Nicholson, K. Liu, H. Chen, Y. Lin, Y. Shu, W. Zhou, C. J. Schuurmans, Q. R. Lu, Neurons derived from NeuroD1-expressing astrocytes transition through transit-amplifying intermediates but lack functional maturity. Sci Adv 11, eadw9296 (2025).

19. N. Marichal, S. Peron, A. Beltran Arranz, C. Galante, F. Franco Scarante, R. Wiffen, C. Schuurmans, M. Karow, S. Gascon, B. Berninger, Reprogramming astroglia into neurons with hallmarks of fast-spiking parvalbumin-positive interneurons by phospho-site-deficient Ascl1. Sci Adv 10, eadl5935 (2024).

20. A. Herrero-Navarro, L. Puche-Aroca, V. Moreno-Juan, A. Sempere-Ferrandez, A. Espinosa, R. Susin, L. Torres-Masjoan, E. Leyva-Diaz, M. Karow, M. Figueres-Onate, L. Lopez-Mascaraque, J. P. Lopez-Atalaya, B. Berninger, G. Lopez-Bendito, Astrocytes and neurons share region-specific transcriptional signatures that confer regional identity to neuronal reprogramming. Sci Adv 7, (2021).

21. C. Lentini, M. d’Orange, N. Marichal, M. M. Trottmann, R. Vignoles, L. Foucault, C. Verrier, C. Massera, O. Raineteau, K. K. Conzelmann, S. Rival-Gervier, A. Depaulis, B. Berninger, C. Heinrich, Reprogramming reactive glia into interneurons reduces chronic seizure activity in a mouse model of mesial temporal lobe epilepsy. Cell Stem Cell 28, 2104–2121 e2110 (2021).

22. W. Tai, W. Wu, L. L. Wang, H. Ni, C. Chen, J. Yang, T. Zang, Y. Zou, X. M. Xu, C. L. Zhang, In vivo reprogramming of NG2 glia enables adult neurogenesis and functional recovery following spinal cord injury. Cell Stem Cell 28, 923–937 e924 (2021).

23. W. Niu, T. Zang, Y. Zou, S. Fang, D. K. Smith, R. Bachoo, C. L. Zhang, In vivo reprogramming of astrocytes to neuroblasts in the adult brain. Nat Cell Biol 15, 1164–1175 (2013).

24. N. L. Jorstad, M. S. Wilken, W. N. Grimes, S. G. Wohl, L. S. VandenBosch, T. Yoshimatsu, R. O. Wong, F. Rieke, T. A. Reh, Stimulation of functional neuronal regeneration from Muller glia in adult mice. Nature 548, 103–107 (2017).

25. O. Torper, U. Pfisterer, D. A. Wolf, M. Pereira, S. Lau, J. Jakobsson, A. Bjorklund, S. Grealish, M. Parmar, Generation of induced neurons via direct conversion in vivo. Proc Natl Acad Sci U S A 110, 7038–7043 (2013).

26. C. Boudreau-Pinsonneault, L. A. David, J. A. Lourenco Fernandes, A. Javed, M. Fries, P. Mattar, M. Cayouette, Direct neuronal reprogramming by temporal identity factors. Proc Natl Acad Sci U S A 120, e2122168120 (2023).

27. L. L. Wang, C. Serrano, X. Zhong, S. Ma, Y. Zou, C. L. Zhang, Revisiting astrocyte to neuron conversion with lineage tracing in vivo. Cell 184, 5465–5481 e5416 (2021).

28. T. Hoang, D. W. Kim, H. Appel, M. Ozawa, S. Zheng, J. Kim, S. Blackshaw, Ptbp1 deletion does not induce astrocyte-to-neuron conversion. Nature 618, E1–E7 (2023).

29. D. Leib, Y. H. Chen, A. M. Monteys, B. L. Davidson, Limited astrocyte-to-neuron conversion in the mouse brain using NeuroD1 overexpression. Mol Ther 30, 982–986 (2022).

30. R. Bocchi, G. Masserdotti, M. Gotz, Direct neuronal reprogramming: Fast forward from new concepts toward therapeutic approaches. Neuron 110, 366–393 (2022).

31. F. R. Ali, K. Cheng, P. Kirwan, S. Metcalfe, F. J. Livesey, R. A. Barker, A. Philpott, The phosphorylation status of Ascl1 is a key determinant of neuronal differentiation and maturation in vivo and in vitro. Development 141, 2216–2224 (2014).

32. S. Li, P. Mattar, R. Dixit, S. O. Lawn, G. Wilkinson, C. Kinch, D. Eisenstat, D. M. Kurrasch, J. A. Chan, C. Schuurmans, RAS/ERK signaling controls proneural genetic programs in cortical development and gliomagenesis. J Neurosci 34, 2169–2190 (2014).

33. L. Lim, D. Mi, A. Llorca, O. Marin, Development and Functional Diversification of Cortical Interneurons. Neuron 100, 294–313 (2018).

34. J. E. Long, I. Cobos, G. B. Potter, J. L. Rubenstein, Dlx1&2 and Mash1 transcription factors control MGE and CGE patterning and differentiation through parallel and overlapping pathways. Cereb Cortex 19 **Suppl 1**, i96–106 (2009).

35. D. G. Miller, M. A. Adam, A. D. Miller, Gene transfer by retrovirus vectors occurs only in cells that are actively replicating at the time of infection. Mol Cell Biol 10, 4239–4242 (1990).

36. R. A. Katz, J. G. Greger, A. M. Skalka, Effects of cell cycle status on early events in retroviral replication. J Cell Biochem 94, 880–889 (2005).

37. C. Galante, N. Marichal, F. F. Scarante, L. M. Ghayad, Y. Shi, C. Schuurmans, B. Berninger, S. Peron, Enhanced proliferation of oligodendrocyte progenitor cells following retrovirus mediated Achaete-scute complex-like 1 overexpression in the postnatal cerebral cortex in vivo. Front Neurosci 16, 919462 (2022).

38. S. A. Anderson, D. D. Eisenstat, L. Shi, J. L. Rubenstein, Interneuron migration from basal forebrain to neocortex: dependence on Dlx genes. Science 278, 474–476 (1997).

39. I. Toudji, A. Toumi, E. Chamberland, E. Rossignol, Interneuron odyssey: molecular mechanisms of tangential migration. Front Neural Circuits 17, 1256455 (2023).

40. Y. H. Liu, J. W. Tsai, J. L. Chen, W. S. Yang, P. C. Chang, P. L. Cheng, D. L. Turner, Y. Yanagawa, T. W. Wang, J. Y. Yu, Ascl1 promotes tangential migration and confines migratory routes by induction of Ephb2 in the telencephalon. Sci Rep 7, 42895 (2017).

41. N. C. Spitzer, Electrical activity in early neuronal development. Nature 444, 707–712 (2006).

42. S. S. Rosenberg, N. C. Spitzer, Calcium signaling in neuronal development. Cold Spring Harb Perspect Biol 3, a004259 (2011).

43. B. Wamsley, G. Fishell, Genetic and activity-dependent mechanisms underlying interneuron diversity. Nat Rev Neurosci 18, 299–309 (2017).

44. L. Felix, J. Stephan, C. R. Rose, Astrocytes of the early postnatal brain. Eur J Neurosci 54, 5649–5672 (2021).

45. A. Semyanov, C. Henneberger, A. Agarwal, Making sense of astrocytic calcium signals - from acquisition to interpretation. Nat Rev Neurosci 21, 551–564 (2020).

46. A. Cooper, A. G. Mora, G. Herrera-Oropeza, A. B. Arranz, N. Marichal, Y. Shi, S. Leaman, K. L. Gonzalez, M. Strom, H. Fursham, F. Guillemot, B. Berninger, Dlx2 reprograms the transcriptome and laminar position of glia-derived Ascl1-induced interneurons. bioRxiv, 2026.2001.2002.697435 (2026).

47. N. V. De Marco García, T. Karayannis, G. Fishell, Neuronal activity is required for the development of specific cortical interneuron subtypes. Nature 472, 351–355 (2011).

48. D. Linaro, B. Vermaercke, R. Iwata, A. Ramaswamy, B. Libe-Philippot, L. Boubakar, B. A. Davis, K. Wierda, K. Davie, S. Poovathingal, P. A. Penttila, A. Bilheu, L. De Bruyne, D. Gall, K. K. Conzelmann, V. Bonin, P. Vanderhaeghen, Xenotransplanted Human Cortical Neurons Reveal Species-Specific Development and Functional Integration into Mouse Visual Circuits. Neuron 104, 972–986 e976 (2019).

49. S. Falkner, S. Grade, L. Dimou, K. K. Conzelmann, T. Bonhoeffer, M. Gotz, M. Hubener, Transplanted embryonic neurons integrate into adult neocortical circuits. Nature 539, 248–253 (2016).

50. S. Goebbels, I. Bormuth, U. Bode, O. Hermanson, M. H. Schwab, K. A. Nave, Genetic targeting of principal neurons in neocortex and hippocampus of NEX-Cre mice. Genesis 44, 611–621 (2006).

51. C. Stringer, T. Wang, M. Michaelos, M. Pachitariu, Cellpose: a generalist algorithm for cellular segmentation. Nat Methods 18, 100–106 (2021).

52. M. Pachitariu, C. Stringer, M. Dipoppa, S. Schröder, L. F. Rossi, H. Dalgleish, M. Carandini, K. D. Harris, Suite2p: beyond 10,000 neurons with standard two-photon microscopy. (2017).

53. P. Thévenaz, U. E. Ruttimann, M. Unser, A pyramid approach to subpixel registration based on intensity. IEEE Trans Image Process 7, 27–41 (1998).

54. C. T. Rueden, J. Schindelin, M. C. Hiner, B. E. DeZonia, A. E. Walter, E. T. Arena, K. W. Eliceiri, ImageJ2: ImageJ for the next generation of scientific image data. BMC Bioinformatics 18, 529 (2017).

55. M. Arzt, J. Deschamps, C. Schmied, T. Pietzsch, D. Schmidt, P. Tomancak, R. Haase, F. Jug, LABKIT: Labeling and Segmentation Toolkit for Big Image Data. Frontiers in Computer Science 4, (2022).

56. S. Seabold, J. Perktold, in Proceedings of the 9th Python in Science Conference, S. e. v. d. Walt, J. Millman, Eds. (2010), pp. 92–96.

57. J. Ho, T. Tumkaya, S. Aryal, H. Choi, A. Claridge-Chang, Moving beyond P values: data analysis with estimation graphics. Nat Methods 16, 565–566 (2019).

